# Perforin-2 is a pore-forming effector of endocytic escape in cross-presenting dendritic cells

**DOI:** 10.1101/2023.01.31.525875

**Authors:** Pablo Rodríguez-Silvestre, Marco Laub, Alexandra K. Davies, Julia P. Schessner, Patrycja A. Krawczyk, Benjamin J. Tuck, William A. McEwan, Georg H.H. Borner, Patrycja Kozik

**Author notes:** Corresponding author: Patrycja Kozik.

## Abstract

During initiation of antiviral and antitumour T cell-mediated immune responses, dendritic cells (DCs) cross-present exogenous antigens on MHC class I. Cross-presentation relies on the unique ‘leakiness’ of endocytic compartments in DCs, whereby internalised proteins escape into the cytosol for proteasome-mediated generation of MHC I-binding peptides. Given that type 1 conventional DCs excel at cross-presentation, we searched for cell-type specific effectors of endocytic escape. We devised an escape assay suitable for genetic screening and identified a pore-forming protein, perforin-2, as a dedicated effector exclusive to cross-presenting cells. Perforin-2 is recruited to antigen-containing compartments, where it undergoes maturation, releasing its pore-forming domain. *Mpeg1*^-/-^ mice fail to efficiently prime CD8^+^ T cells to cell-associated antigens, revealing an important role of perforin-2 in cytosolic entry of antigens during cross-presentation.

**One-Sentence Summary:** Pore-forming protein perforin-2 is a dedicated effector of endocytic escape specific to cross-presenting cells

## Main text

The integrity of endosomal and lysosomal membranes is critical to protect the cell against extracellular pathogens and toxins, as well as from the activity of lysosomal hydrolases. Yet dendritic cells (DCs) allow internalised proteins to be delivered from endocytic organelles into the cytosol where they can be proteolytically processed for presentation on MHC class I molecules (*1*). The ability of dendritic cells to present exogenous peptides on endogenous MHC class I is termed cross-presentation and is critical for initiation of cytotoxic T cell (CTL) immune responses to antigens not expressed in DCs such as tumour neoantigens or antigens derived from virally infected cells (*2*–*4*).

Cross-presentation via the endosome-to-cytosol pathway was first reported over 20 years ago (*5*). Since then, various mechanisms have been shown to facilitate endocytic escape and promote cross-presentation (*6*–*10*). Early studies proposed that escape is mediated by protein channels (such as Sec61), which are recruited from the endoplasmic reticulum to endosomes in cross-presenting cells (*7*). More recent data suggest that escape occurs through unrepaired damage to endosomal membranes (e.g., due to ROS-driven lipid peroxidation) (*6*, *9*, *10*). Both models imply that the unusual ‘leakiness’ of endocytic compartments in cross-presenting DCs might not rely on any cell type-specific effectors, but rather on the regulation of ubiquitously expressed proteins through signalling (*11*, *12*) and trafficking (*13*) events unique to cross-presenting cells.

To gain a better understanding of how cross-presenting cells control endocytic escape, we set out to perform a genetic screen for DC-specific regulators. To this end, we developed a flow cytometry-based strategy to monitor endocytic escape in individual cells and adopted it for a CRISPR/Cas9-based screen in a cell type specialised in cross-presentation, conventional dendritic cells 1 (cDC1).

### Saporin-puromycin assay to monitor the efficiency of antigen import

To monitor endocytic escape in cross-presenting DCs, we used a type I ribosome inactivating protein (RIP) saporin (Fig. 1A) (*5*). Once in the cytosol, RIPs cause rapid translation arrest through depurination of the sarcin-ricin loop in the 28 subunit of the ribosome (*14*). While type II RIPs, such as ricin, comprise a domain that facilities entry into the cytosol, cytosolic delivery of type I RIPs is dependent on the cell intrinsic efficiency of endosome-to-cytosol transport (*5*). To detect saporin-induced translation inhibition, we used puromycin, a structural analogue of aminoacyl tRNAs, which is incorporated into nascent polypeptides at a rate that correlates with translation efficiency (*15*–*17*). Intracellular puromycylated polypeptides can be labelled with a fluorescent monoclonal antibody to puromycin (12D10) allowing for flow cytometry-based readout of translation efficiency and thereby of endocytic escape (Fig. 1A).

**Fig 1.**
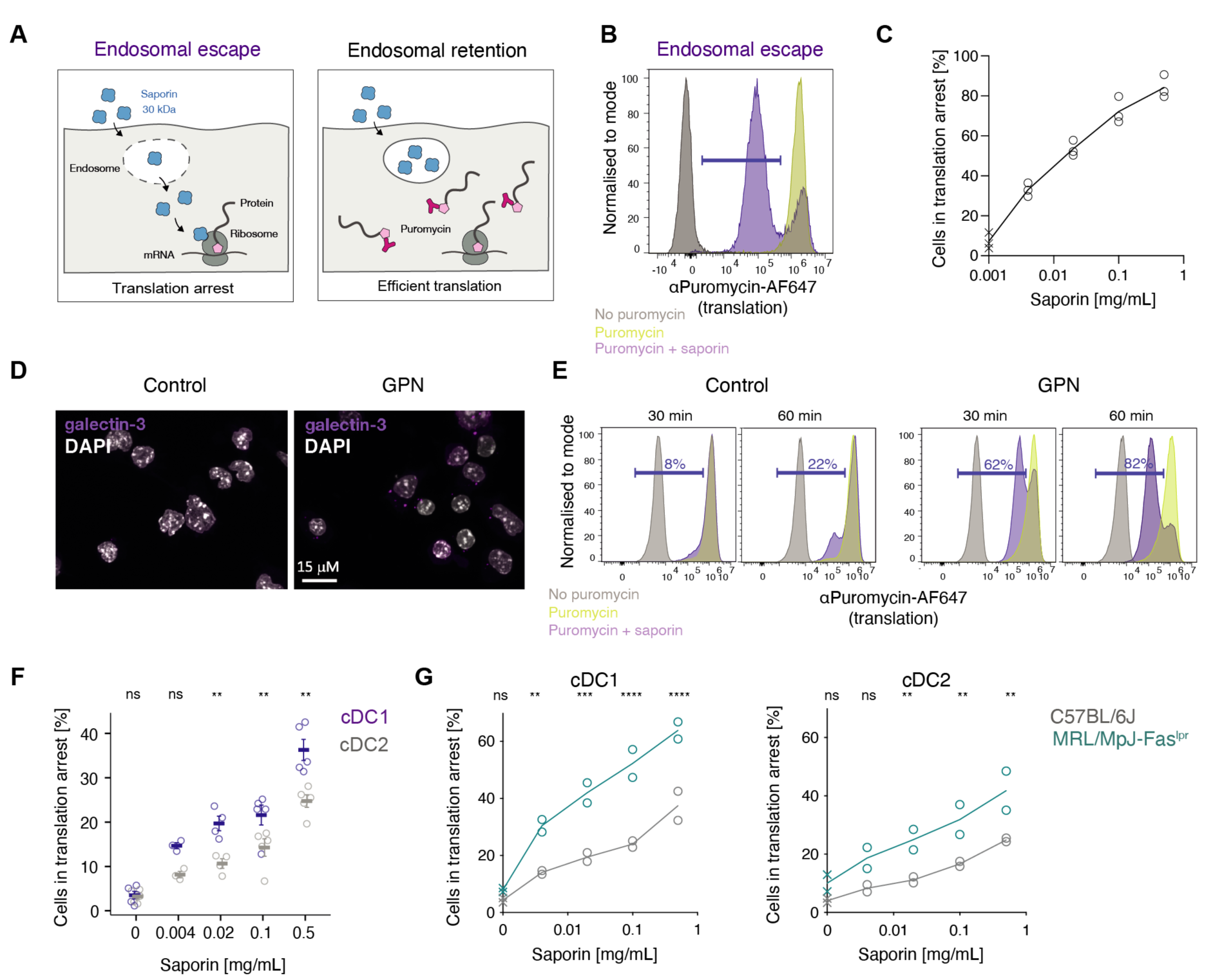
Saporin-puromycin assay to monitor endocytic escape in DCs. **(A)** Schematic representation of the saporin-puromycin assay. Cells are pulsed with saporin for 2 h and translation is then monitored by labelling nascent polypeptides with puromycin for 30 min. Incorporated puromycin can then be detected with an αPuromycin Ab and flow cytometry. If saporin is retained within the endosomes, translation remains high. When saporin escapes into the cytosol it depurinates ribosomes inducing translation arrest. **(B)** Representative flow cytometry plot from the saporin-puromycin assay described in (A). MutuDCs were incubated with 0.5 mg/ml of saporin followed by 0.01 mg/mL puromycin (purple histogram), with puromycin alone (yellow) or in media only (grey). Cells in translation arrest are denoted by the purple gate. **(C)** MutuDCs were incubated with different amounts of saporin as in (B) and the percentage of puromycin^low^ cells quantified by flow cytometry. Data represent three independent experiments. For gating strategy see fig. S1B. **(D)** Confocal microscopy images of MutuDCs treated with 33 μM GPN for 10 min. Cells were stained with DAPI (white) and for galectin-3 (magenta). **(E)** MutuDCs were incubated with 0.5 mg/mL saporin for the indicated time in the presence or absence of 33 μM GPN. Following a 30 min puromycin chase, translation inhibition was monitored by flow cytometry. Data represent three independent experiments. **(F)** CD11c+ enriched splenic DCs from C57BL/6J were incubated with saporin for 2 h, followed by a 30 min puromycin chase. Translation inhibition was monitored by flow cytometry. Data represent mean and SEM for five independent experiments, ns, not significant; **, P<0.01 using a multiple unpaired t-test (two-stage step-up, Benjamini, Krieger and Yekutieli). For gating strategy see fig. S1C. **(G)** CD11c^+^ enriched splenic DCs from C57BL/6J or MRL/MpJ-Fas^lpr^ were incubated with saporin for 2 h, followed by a 30 min puromycin chase. Translation inhibition was monitored by flow cytometry. Data represent mean and SEM for two independent experiments each with two technical replicates, ns, not significant; \**P*<0.5; ** *P*<0.01; *** *P*<0.001; \*\*\*\**P*<0.0001 using a multiple unpaired t-test (alpha = 0.5). For gating strategy see fig. S1C and fig. S1D.

cDC1s, a subset of DCs that excels in cross-presentation *in vivo*(*18*) have been reported to have the most efficient endocytic escape pathway (*18*, *19*). Therefore, we set up the assay using a murinecDC1-like cell line, MutuDCs (*12*, *20*, *21*). A 30 min pulse with puromycin led to its efficient incorporation which could be blocked by a 2 h pretreatment with the translation inhibitor cycloheximide confirming that puromycylation in MutuDCs is sensitive to changes in the rate of translation (fig. S1A). Next, we pulsed the cells with saporin and observed an increase in the number of cells with low puromycin signal indicating translation arrest due to saporin escape (Fig. 1B). The saporin-induced translational arrest was dose-dependent (Fig. 1C), confirming that our assay can accurately measure the efficiency of endocytic escape.

To demonstrate that endocytic escape of saporin is the rate-limiting step in the saporin-puromycin assay, we disrupted the integrity of endocytic compartments with glycyl-L-phenylalanine 2-naphthylamide (GPN). GPN is a cathepsin C substrate that induces osmotic lysis of lysosomes leading to release of lysosomal contents (*22*). In MutuDCs, 33 μM GPN caused endocytic damage visualised by staining for galectin 3, a marker of damaged compartments (*23*) (Fig. 1D). In the saporin-puromycin assay, the population of cells with arrested translation appeared with faster kinetics in the presence of GPN, confirming endocytic escape of saporin is rate-limiting in the saporin-puromycin assay (Fig. 1E).

### The saporin-puromycin assay recapitulates physiologically relevant differences in the efficiency of endocytic escape

To test whether the saporin-puromycin assay can capture physiological differences in the efficiency of endocytic escape, we first focused on primary murine DCs subsets, cDC1s and cDC2s (previously reported as less efficient in antigen import and cross-presentation than cDC1s (*19*)). We performed the saporin-puromycin assay *ex vivo,* on splenic cDCs isolated by negative selection, and, in agreement with previous data, we demonstrated that saporin escape was more efficient in cDC1s (Fig. 1F).

We also analysed the efficiency of saporin escape in DCs from lupus-prone mice, homozygous for the lymphoproliferation spontaneous mutation (MRL/MpJ-Fas^lpr^/J). Macrophages from the MRL mice have a defect in lysosomal stability resulting in leakage of lysosomal contents and excessive activation of cytosolic sensors (*24*). We performed the saporin-puromycin assay on CD11c ^+^ pre-enriched splenic DCs and found that saporin entered the cytosol more efficiently in cDC1s (and cDC2s) from the MRL mice compared to wild type C57BL/6J (Fig. 1G). Together, these experiments demonstrate that the saporin-puromycin assay recapitulates previously observed differences in the efficiency of endocytic escape.

### A CRISPR/Cas9 screen for regulators of endocytic escape

To identify regulators of endocytic escape in DCs, we employed the saporin-puromycin assay in a CRISPR/Cas9 genetic screen. Considering the difference in efficiency of endocytic escape between the cDC1 and cDC2s (*25*) (Fig. 1F), we focused on 281 genes highly expressed in cDC1s (Fig. 2A). We generated a lentiviral mini-library using a BFP-containing plasmid, where each of the upregulated genes was targeted with four sgRNAs (Fig. 2B). We infected 4x10^6^ MutuDCs stably expressing Cas9, sorted for BFP^+^ cells four days later, and expanded the cell library for a minimum of 10 days. We confirmed that all sgRNAs were recovered from MutuDCs when compared to the original plasmid library (table S1), except for sgRNAs targeting *Irf8*, which were significantly depleted (Fig. 2C). Considering Irf8 is a transcription factor required for cDC1 differentiation and for survival of terminally differentiated cDC1s (*26*), this analysis confirmed the suitability of MutuDCs as a cDC1 model system and demonstrated that the efficiency of gene editing in MutuDCs is sufficient for genetic screening.

**Fig 2.**
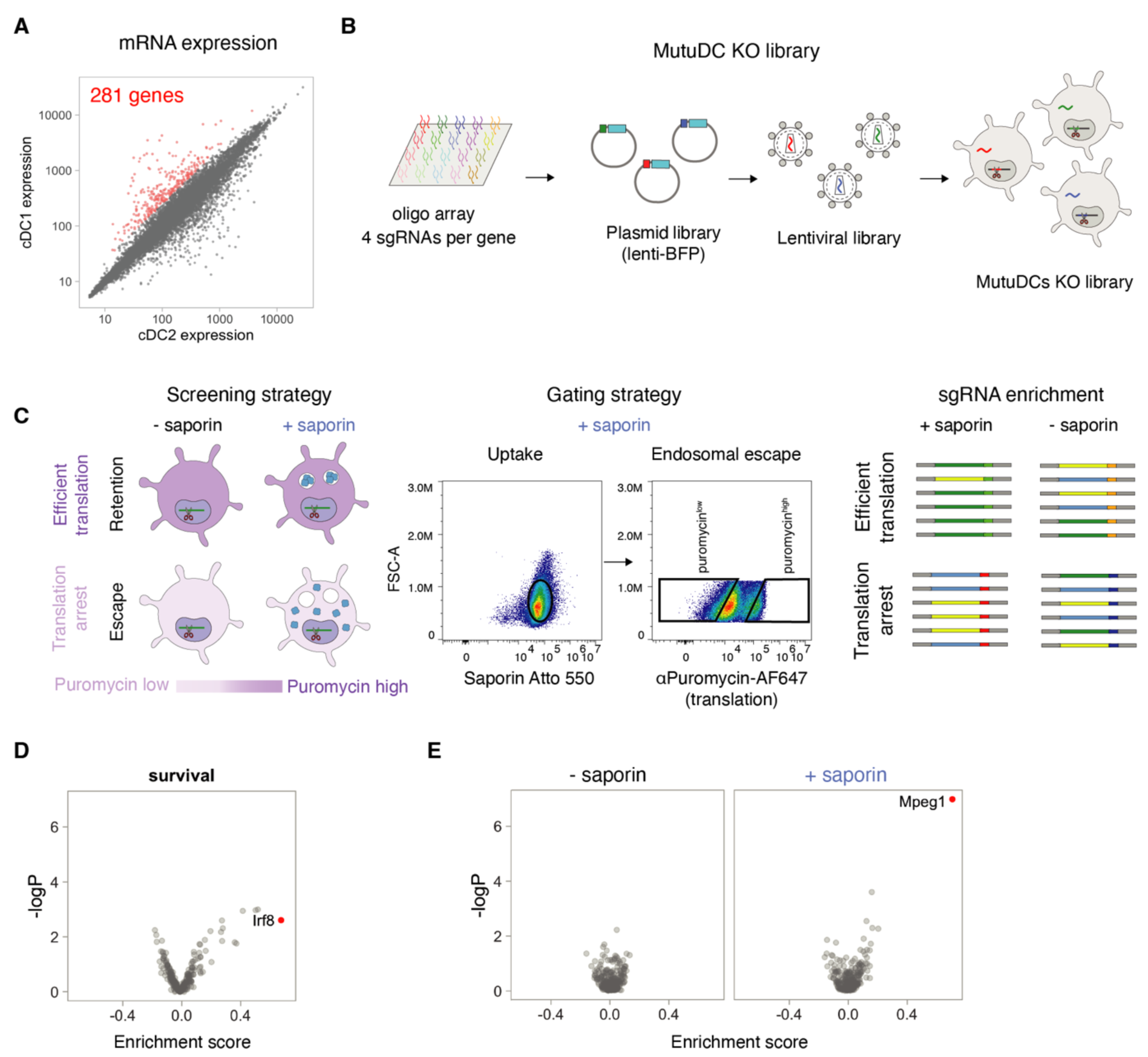
Genetic screen identifies Mpeg1 as a candidate regulator of endocytic escape in DCs. **(A)** cDC1 and cDC2 transcript comparison (data from the ImmGen consortium). 281 genes (coloured in red) met the log2(cDC1/cDC2) > 1.3 cut-off and were included in the CRIPSR/Cas9 library. **(B)** The CRISPR/Cas9 library consisted of 4 sgRNAs per gene in a BFP-expressing lentiviral vector. Cas9-expressing MutuDCs were transduced with at an MOI of 0.3. Cells were sorted for BFP expression and expanded. **(C)** Schematic representation of the CRISPR/Cas9 screen. For the screen, the saporin-puromycin assay was performed with a 2 h pulse of 0.5 mg/mL saporin (1:11 ratio of saporin Atto550 to unlabelled). Cells were first gated on the basis of the Atto550 MFI, and cells with either high (saporin escape) or low (saporin retention) puromycin labelling were collected. Puromycin high and low cells were also collected in the absence of saporin to identify guides that might have a global effect on translation. Genomic DNA was then isolated and prepared for next generation sequencing of the sgRNAs. **(D)** Volcano plot showing the sgRNAs enrichment analysis for the MutuDC library relative to the starting plasmid library. Each of the dots represents one targeted gene. **(E)** Volcano plots showing the sgRNAs enrichment analysis for the indicated screens. Each of the dots represents one targeted gene. Data represent the combined mean enrichment scores and the non-adjusted p values from three independent experiments (Fisher’s method).

To identify genes required for endocytic escape, we performed the saporin-puromycin assay using 0.5 mg/mL saporin spiked 11:1 with fluorescently labelled saporin-Atto 550. To ensure that the difference in puromycin incorporation was not due to a difference in uptake efficiency, we only collected the cells that took up similar amounts of saporin (Fig. 2C). We sorted the cells into two bins: puro^low^ (saporin escape and translation arrest) and puro^high^ (saporin retention and efficient translation). To control for possible differences in puromycin incorporation due to saporin-independent inhibition of translation, we also collected puro^low^ and puro^high^ populations from MutuDCs labelled in the absence of saporin. None of the sgRNAs affected the overall translation rate (Fig. 2D). The strongest hit identified in saporin-pulsed cells was *Mpeg1*, with four *Mpeg1*-targetting sgRNAs enriched in puro^low^ vs puro^high^ populations (Fig. 2E and fig. S2A).

### Perforin-2 is necessary for endocytic escape in DCs

*Mpeg1* encodes perforin-2, a member of the membrane attack complex (MAC) and perforin superfamily (MACPF) of pore-forming proteins (*27*). Recent cryo-EM studies demonstrated that, like perforin-1 and complement, perforin-2 can form oligomeric pores on liposomes (*28*, 1. *29*) (Fig. 3A). It has been initially proposed that the function of the perforin-2 pores is to facilitate killing of intravacuolar bacteria in infected cells (*30*), but these results were not replicated in a more recent study (*31*). Here, we explored whether perforin-2 can function as an effector of endocytic escape in DCs. To this end, we generated a *Mpeg1*^KO^ MutuDC line by combining two sgRNAs targeting *Mpeg1* (expressed from a BFP-containing plasmid) and confirmed protein depletion in sorted BFP^+^ cells by Western Blotting (fig. S2B). To account for the effect of passage numbers on MutuDCs behaviour (*21*), we cultured control cells expressing non-targeting (NT) sgRNA in parallel with the KO line.

**Fig 3.**
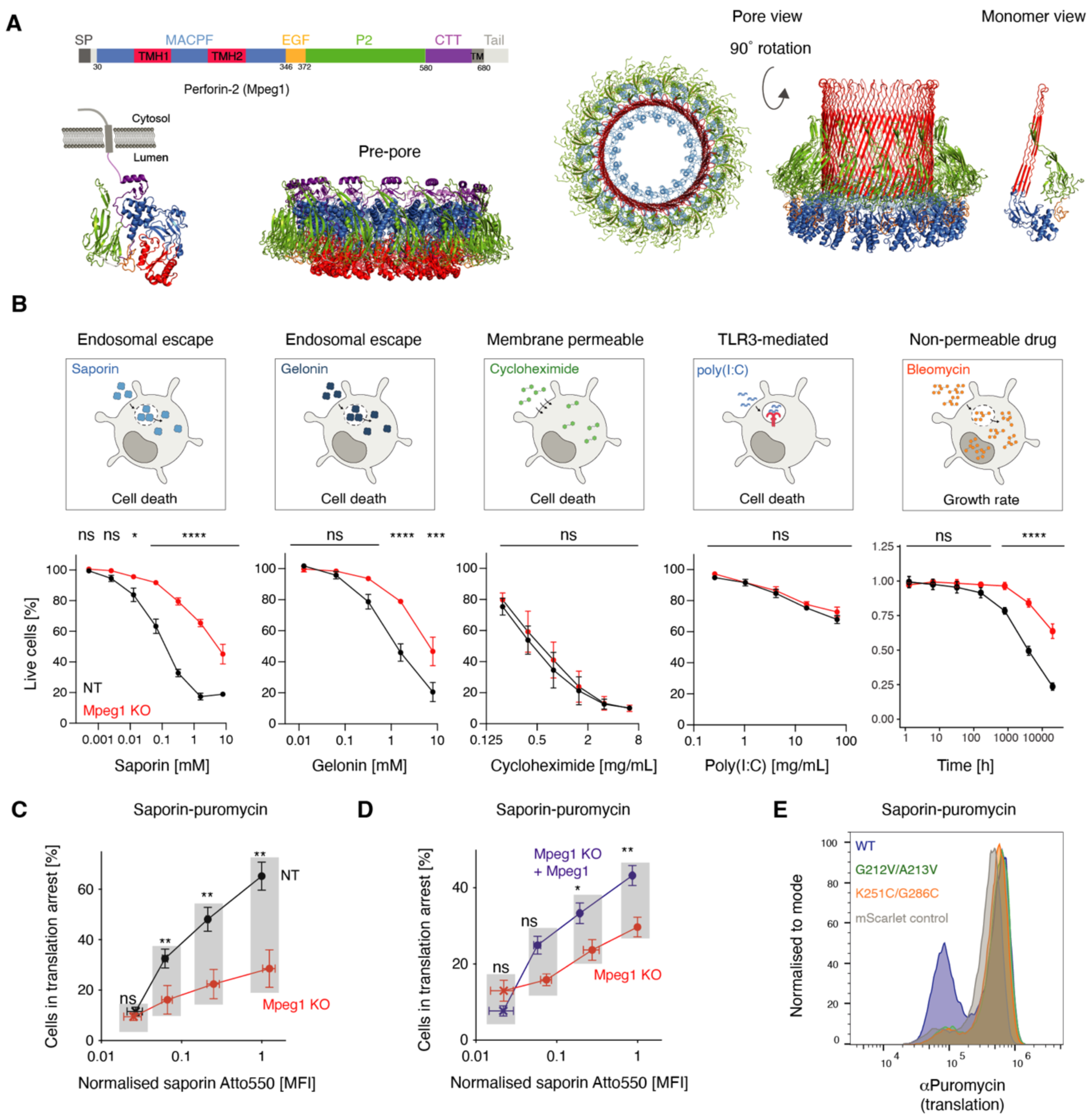
Perforin-2 facilitates endocytic escape of diverse cargo. **(A)** Schematic representation of the different perforin-2 domains alongside structures of single subunit and hexadecameric perforin-2 in pre-pore (PDB ID: 6U2K and PDB ID: 6SB3) and pore-forming conformations (PDB ID: 6SB5). **(B)** *Mpeg1*^KO^ and NT MutuDCs were treated for 24 h with either saporin, gelonin or cycloheximide or for 48 h with HMW-Poly(I:C). Cells were stained with a fixable live/dead stain, and viability was assessed by flow cytometry. For bleomycin, the cells were plated in the presence or absence of bleomycin and cultured in an Incucyte® for 48 h to monitor the growth rate. Data represent mean and SEM of three independent experiments with four technical replicates each, ns, not significant; \**P*<0.5; \*\**P*<0.01; \*\*\**P*<0.001; \*\*\*\**P*<0.0001 using a multiple t-test (Bonferroni-Dunn). **(C and D)** Quantification of translation arrest in (C) *Mpeg1*^KO^ and NT MutuDCs or (D) *Mpeg1*^KO^ and *Mpeg1*-complemented *Mpeg1*^KO^ MutuDCs. Cells were pulsed with saporin (1:11, saporin-Atto 550:unlabelled) for 2 h, and translation was monitored by a 30 min puromycin chase. The X axis represents Atto 550 MFI, normalised to the (C) NT MutuDC or (D) *Mpeg1*^KO^ MutuDCs Atto 550 MFI at the highest saporin concentration. Data represent mean and SEM of three independent experiments, ns, not significant; \**P*<0.5; ** *P*<0.01; *** *P*<0.001; \*\*\*\**P*<0.0001 using a multiple unpaired t-test (two-stage step-up, Benjamini, Krieger and Yekutieli). Significance symbols in the plot refer to the differences in proportion of cells in translation arrest. Differences in saporin Atto 550 MFI were not significant. **(E)** *Mpeg1*^KO^ MutuDCs were reconstituted with the indicated *Mpeg1* mutants or with mScarlet only and used in the saporin-puromycin assay with a 2 h pulse at 0.1 mg/ml saporin. Data are representative for three independent experiments.

Since translation inhibition leads to cell death, we initially evaluated whether the disruption of *Mpeg1* protects the cells from the cytotoxic effects of saporin and other cytosolic toxins. We incubated control and *Mpeg1*^KO^ MutuDCs with saporin for 24 h and quantified surviving cells by flow cytometry. Indeed, Mpeg1^KO^ MutuDCs were more resistant to saporin-mediated cell death, and we observed similar levels of protection from death induced by a different RIP, gelonin (Fig. 3B). Importantly, the KO cells were not protected against cell death initiated by molecules that do not need to access the cytosol or against membrane permeable agents which can enter the cytosol freely. For example, knocking out *Mpeg1* conferred no resistance to Poly (I:C), which in cDC1s induces cell death via endosomal Toll like receptor 3, Tlr3 (*21*) or to cycloheximide, a membrane-permeable translation inhibitor (Fig. 3B). We also tested the sensitivity of Mpeg1^KO^ DCs to a glycopeptide chemotherapeutic, bleomycin A2, which induces DNA damage, but due to its hydrophilicity does not enter the cells efficiently (*32*). We used automated imaging to monitor the growth rate of NT and *Mpeg1*^KO^ MutuDC cultured in the presence of increasing concentrations of bleomycin and observed that loss of perforin-2 renders MutuDCs more resistant to bleomycin-mediated cytotoxicity (Fig. 3B).

We went on to verify that the observed sensitivity of perforin-2-expressing DCs to cytosolic toxins is due to an efficient escape pathway rather than due to inefficient uptake (Fig. 3C). In the saporin-puromycin assay with Atto 550-labelled saporin, endocytic escape was impaired in the *Mpeg1*^KO^ DCs, even though NT and KO cells internalised similar amounts of fluorescent saporin (Fig. 3C). We could also rescue saporin import without affecting uptake by transducing *Mpeg1*^KO^ cells with lentiviruses expressing full length sgRNA-resistant *Mpeg1* (Fig. 3D and fig. S2C).

Finally, we addressed whether the pore-forming ability of perforin-2 is required for endocytic escape. Perforin-2 forms pores by oligomerisation and unwinding of two helices in the MACPF domain into β sheets (indicated in red in Fig. 3A) (*28*, *29*). To test whether this conformational change is required for perforin-2 activity in DCs, we generated two mutants: *Mpeg1*^G212V/A213V^ with mutations in the conserved MACPF motif (*33*) and *Mpeg1*^K251C/G286C^, where we introduced a disulphide bond previously shown to constrain one of the pore-forming helices preventing pore formation *in vitro* (*28*). Both, *Mpeg1*^G212V/A213V^ and *Mpeg1*^K251C/G286C^, were expressed at similar levels to Mpeg1*^WT^*, but neither rescued saporin escape in *Mpeg1*^KO^ MutuDCs (Fig. 3E and fig. S2D). Altogether, these data suggest that perforin-2 pores mediate endocytic escape in cross-presenting DCs.

### Perforin-2 is sufficient for endocytic escape in non-immune cells

Since perforin-2 is restricted to antigen presenting cells (*25*, *34*), we asked whether it is sufficient to drive endocytic escape in non-immune cells. To this end, we generated HEK293Ts and HeLa cells expressing either murine *Mpeg1* together with a fluorescent protein (*mScarlet or BFP),* or the fluorescent protein alone, and used them in a range of endocytic escape assays. Indeed, ectopic expression of perforin-2 was sufficient to promote saporin escape in HEK293Ts (Fig. 4A).

**Fig 4.**
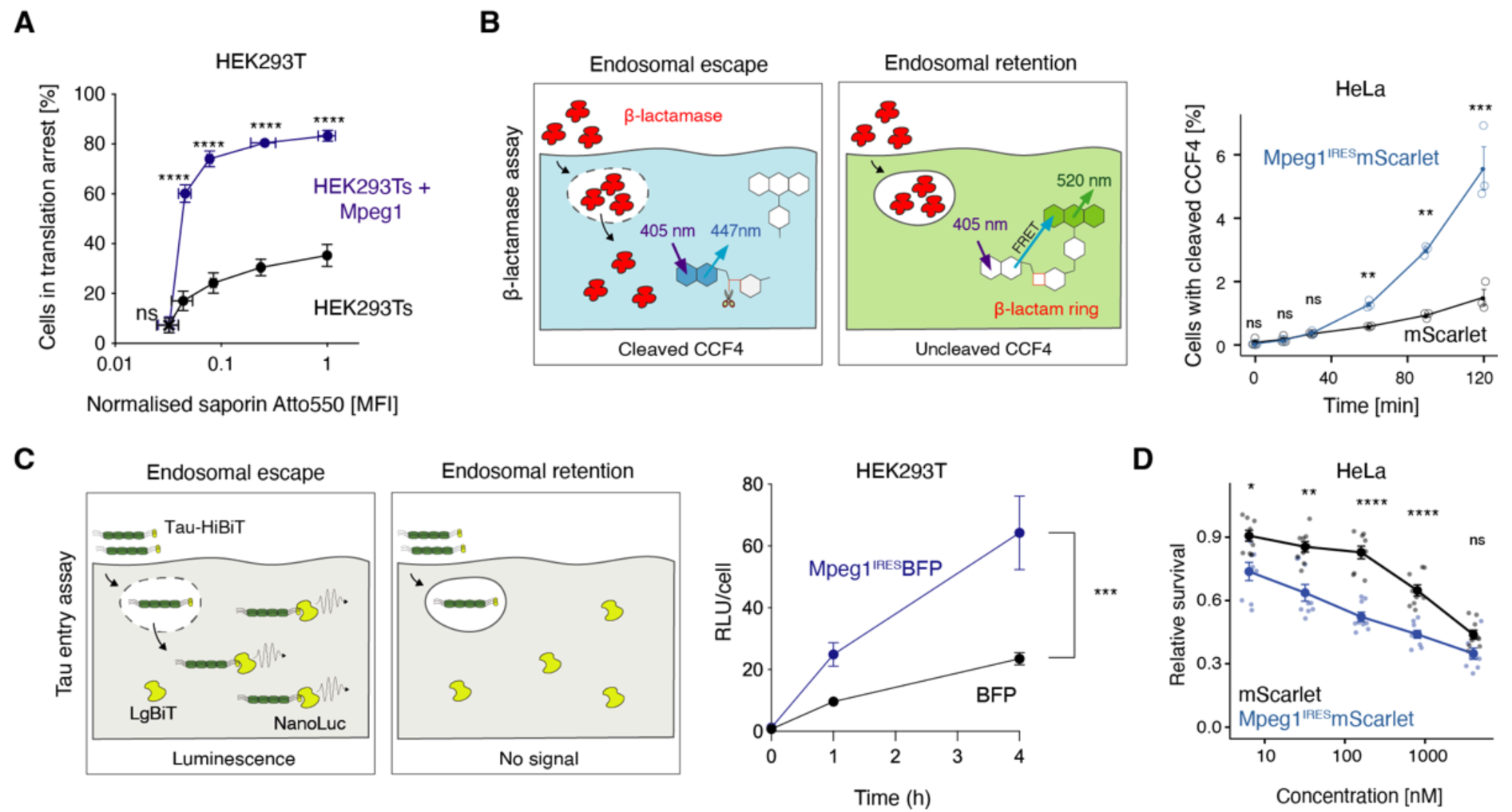
Perforin-2 is sufficient for endocytic escape of cargo in non-immune cells. **(A)** HEK293Ts and Mpeg1-complemented HEK293Ts were pulsed with saporin (1:11, saporin-Atto 550:unlabelled) for 2 h, and translation was monitored by a 30 min puromycin chase. The X axis represents Atto 550 MFI, normalised to the HEK293Ts Atto550 MFI at the highest saporin concentration. Data represent mean and SEM of three independent experiments, ns, not significant; * P<0.5; **P<0.01; ***P<0.001’ ****P<0.0001 using a multiple t-test (Bonferroni-Dunn). Significance symbols on the plot refer to the differences in cells in translation arrest. Any differences in saporin Atto 550 MFI were not significant. **(B)** Cytosolic escape of β-lactamase in cells loaded with the fluorescent β-lactamase substrate CCF4, results in cleavage of the substrate, loss of FRET and emission shift. HeLas expressing either *Mpeg1*^IRES^-mScarlet or mScarlet-only were pulsed with β-lactamase for the indicated time. β-lactamase escape was monitored by measuring the shift in fluorescence emission by flow cytometry. Data represent mean and SEM of three independent experiments, ns, not significant, \**P*<0.5; \*\**P*<0.01; \*\*\**P*<0.001; \*\*\*\**P*<0.0001 using a multiple t-test (two-stage step-up, Benjamini, Krieger and Yekutieli) For gating strategy see fig. S3A. **(C)** To monitor the escape of Tau oligomers, cells expressing NLS-eGFP-LargeBiT were pulsed with Tau-HiBit oligomers. The escape of Tau-HiBiT into the cytoplasm allows binding to the 18 kDA luciferase subunit, LgBiT. This results in reconstitution of catalytic activity and generation of luminescence. NLS-eGFP-LargeBiT HEK293Ts expressing either *Mpeg1*^IRES^-BFP or BFP-only were pulsed with tau-HiBiT for the indicated time. Following substrate addition, luminesce and cell viability were assessed. Relative luminescent units (RLUs) were then normalised to viability per well. Data represent mean and SEM of three independent experiments each with six technical replicates, ***P<0.001 using a paired t-test. **(D)** HEK293Ts were plated in the presence or absence of bleomycin and cultured in an Incucyte® for 48 h to monitor the growth rate. Data represent mean and SEM of three independent experiments with four wells per condition each, ns, not significant; \**P*<0.5; \*\**P*<0.01; \*\*\**P*<0.001; \*\*\*\**P*<0.0001 using a multiple t-test (Bonferroni-Dunn).

We also monitored endocytic escape of β-lactamase using a previously described flow cytometry-based assay (*35*, *36*). Here, the cells are first loaded with a cytosolic dye CCF4 consisting of fluorescein and 7-hydroxycoumarin linked via a β-lactam ring. Escape of β-lactamase into the cytosol results in cleavage of a β-lactam ring and a shift in CCF4 fluorescence from green to blue. We pulsed the cells with β-lactamase for different amounts of time and monitored the proportion of cells with cleaved CCF4 by flow cytometry. Expression of perforin-2 increased the frequency of cells with cleaved CCF4 and thus the efficiency of β-lactamase escape in HeLa cells (Fig. 4B and fig. S3B). We then adopted a recently reported split luciferase assay to monitor endocytic escape of recombinant microtubule-associated protein, tau(*37*). We expressed a large 18 kDa NanoLuc subunit (LgBiT) in the cytosol of control and perforin-2-expressing HEK293Ts and pulsed the cells with oligomers of recombinant tau fused with a high-affinity 11 amino acid-long HiBit peptide. Upon entry into the cytosol, HiBiT binds LgBiT resulting in catalytic activity of luciferase and luminescence in the presence of the enzymatic substrate, Nano-Glo^(R)^. Again, in perforin-2-expressing HEK293Ts, tau-HiBit escape was more efficient compared to the control line (Fig. 4C). Finally, we used the bleomycin-sensitivity assay described above to demonstrate that expression of perforin-2 renders HeLa cells more sensitive to bleomycin-mediated toxicity (Fig. 4D). Collectively, these data show that ectopic expression of perforin-2 is sufficient to drive endocytic escape in non-immune cells.

### Proteolytic cleavage releases the pore-forming domain of perforin-2

Given the cytotoxic potential of pore-forming proteins, we asked how cDC1s regulate pore-formation and restrict it to antigen containing compartments (36, 37).

Perforin-2 is the only known mammalian pore-forming protein with an additional transmembrane domain (TMD) not involved in pore formation (see Fig. 3A). The TMD has been proposed to act as an anchor preventing damage to endogenous membranes and orientating the pore towards intravacuolar bacteria residing in phagosomes (27, 28). We hypothesised that perforin-2 is activated by proteolytic release of the ectodomain in late endocytic compartments. This hypothesis initially emerged from analysis of SILAC-based organellar mapping data which we generated in a previous study (*20*). The organellar maps were prepared by mass spectrometry-based analysis of the fractionated post-nuclear supernatants from control cells and cells treated with drugs that promoted lysosomal leakiness, Prazosin and Tamoxifen (fig. S4) (*38*). In the original maps, perforin-2 had a profile similar to lysosomal proteins. Here, instead of analysing protein profiles, we analysed each tryptic peptide individually, only including endo-or lysosomal proteins (Fig. 5A). We confirmed clustering of individual peptides derived from endosome-or lysosome-resident proteins and of peptides derived from transmembrane or luminal proteins in the lysosome cluster. 15 out of 17 perforin-2-derived peptides clustered with soluble rather than transmembrane lysosomal proteins suggesting that the perforin-2 ectodomain is released from the TMD anchor. The remaining peptides, p340-372 (within the EGF domain) and p629-635 (within a region proximal to the TMD), co-clustered with endosomal proteins (such as Vps35, Fig. 5A) suggesting that they are present in the full-length protein transiting through endosomes but are absent (cleaved) once perforin-2 reaches acidic compartments.

**Fig 5.**
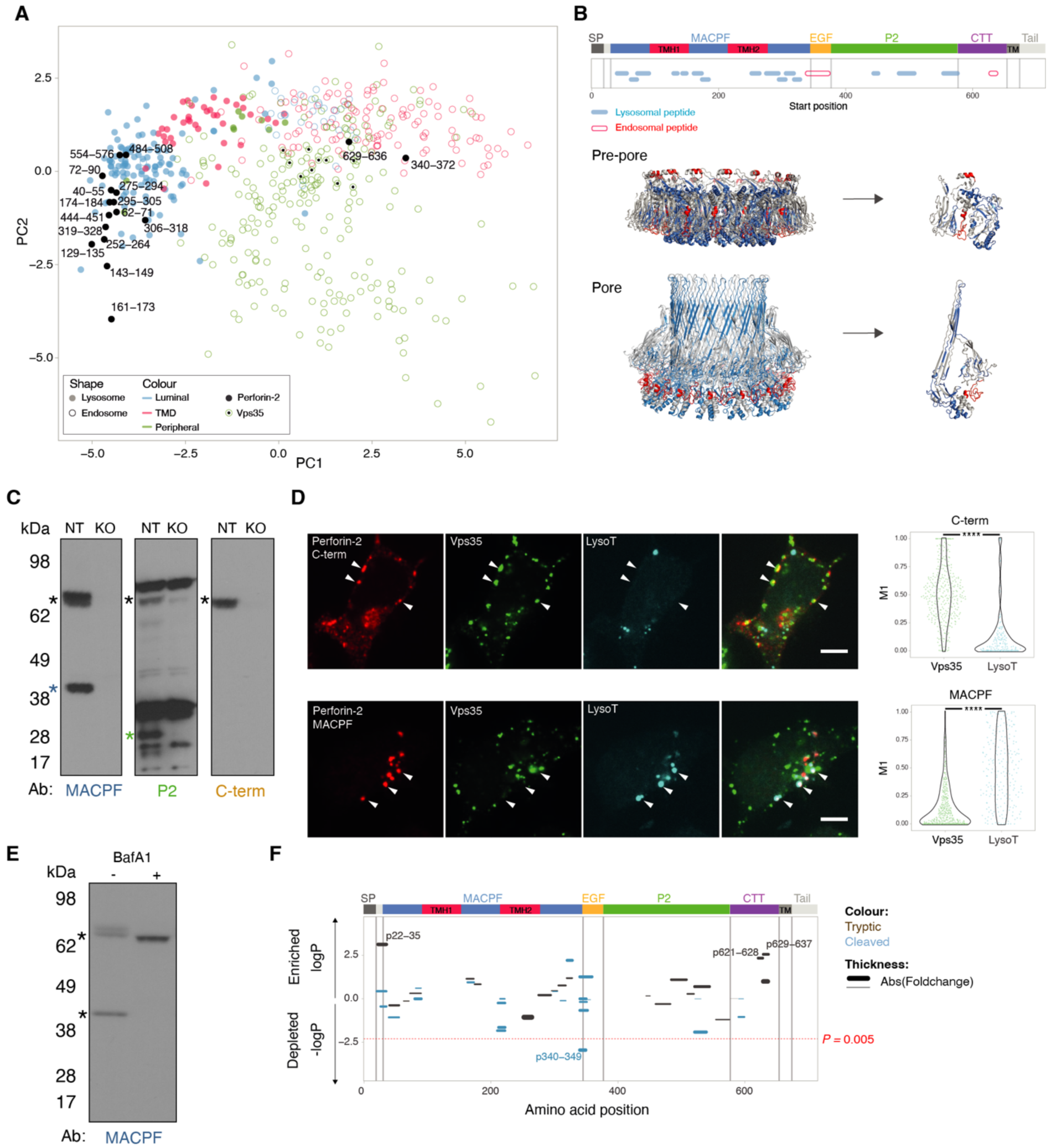
Perforin-2 undergoes proteolytic cleavage releasing its pore forming domain into the organellar lumen. **(A)** Principal component analysis of mass spectrometry-based organellar mapping of MutuDCs (*20*). Lysosomal proteins are represented by filled circles, while endosomal ones are indicated by empty ones. The different colours indicate localisation of the protein within the corresponding organelle. Perforin-2 peptides are displayed as filled black circles regardless of their localisation. **(B)** Mapping of the different lysosomal and endosomal perforin-2 peptides detected by organellar mass spectrometry onto the different perforin-2 domains and structure. **(C)** Perforin-2 levels in *Mpeg1*^KO^ and NT MutuDCs were assessed by Western blot under non-reducing conditions using several antibodies which recognise different perforin-2 epitopes. For ponceau staining see fig. S2E. **(D)** Confocal microscopy images of MutuDCs stained for perforin-2 with either αC-terminal tail or αMACPF antibodies (red), Vps35 (green) and lysotracker (cyan). Data represent two independent experiments with at least 80 cells each. Statistical analysis was performed with a Kolmogorov-Smirnov test.***P < 0.0001. **(E)** Perforin-2 levels in NT MutuDCs treated with 0.5 μM BafA1 for 3 h were assessed by Western blot under reducing conditions using the αMACPF antibody. **(F)** BafA1 induced changes in the abundance of tryptic and cleaved perforin-2 peptides by full proteome mass spectrometry. Control and BafA1 (1 μM) treated cells were analysed by mass spectrometry and peptide abundance was normalised per condition. Fold change in abundance of peptides in BafA1 vs control is shown and the amino acid position indicates the location of the peptides along the different perforin-2 domains. Two-sided student’s t-test was used for statistical analysis. Multiple hypothesis correction was done by permutation based FDR with s0=0.1 using Perseus (table S2).

To confirm that perforin-2 is cleaved near the TMD and within the EGF domain, we probed MutuDC lysates with domain-specific antibodies (Fig. 5C). All of the antibodies (αMACPF, αP2 and the αC-terminal tail) detected full length perforin-2 at 70 kDa. We also detected a 40 kDa band with the αMACPF Ab and a 25 kDa band with the αP2 Ab consistent with the cleavage in the EGF domain. The antibody against the C-terminal tail detected only full-length protein, suggesting that the tail (and most likely the TMD) are rapidly degraded following release of the ectodomain. To test whether this proteolytic processing takes place in lysosomes, we immunostained MutuDCs with the αC-term antibody, and identified full-length perforin-2 in a Vps35+ compartment, but not in Lysotracker-labelled lysosomes (Fig. 5D). In contrast, the αMACPF antibody colocalised with lysotracker, but not with Vps35, suggesting that either the Ab recognises primarily mature perforin-2 or that the majority of the protein is in lysosomes at steady-state. Together, these data suggested that full length perforin-2 resides in (or transits through) endosomes, and upon reaching low pH compartments, it undergoes maturation involving two cleavage events.

To test whether perforin-2 maturation is indeed pH dependent, we used Bafilomycin A1 (BafA1), a V-ATPase inhibitor that interferes with lysosomal acidification (*39*). In BafA1-treated MutuDCs, perforin-2 accumulated as a 70 kDa protein while the 40 kDa MACPF fragment was no longer detected (Fig. 5E). To validate the position of the two putative cleavage sites, we performed a comparative proteomics analysis of tryptic and semi-tryptic perforin-2 peptides from control and BafA1-treated cells (Fig. 5F and table S2). Semi-tryptic peptides are considered to be derived from proteins that were cleaved in the cell, prior to sample processing. Semi-tryptic peptide p340-349 was significantly depleted in BafA1-treated cells (Fig. 5F) consistent with a pH-dependent cleavage between the MACPF and P2. We were also able to reconstitute this cleavage event *in vitro* with asparagine endopeptidase, AEP (fig. S5A and B). Furthermore, tryptic peptides p621-628 and p629-637 were significantly enriched in BafA1-treated cells consistent with the model where TMD-containing C-terminal portion of perforin-2 is lost upon reaching acidic compartments.

In summary, these experiments reveal that perforin-2 activity is controlled by at least two cleavage events. The cleavage in the TMD-proximal CTT domain releases the ectodomain into the lysosomal lumen and is necessary to orient the pore-forming domain towards the endogenous membranes. The cleavage in the EGF domain is not required for pore formation *in vitr*o (*28*, *40*), but may provide additional flexibility to complete pore insertion *in vivo* or may serve to inactivate the pores.

### Perforin-2 undergoes processing in antigen-containing compartments

To demonstrate that perforin-2 undergoes maturation in antigen-containing compartments, we analysed perforin-2 recruitment to phagosomes by phagoFACS (*41*). MutuDCs were pulsed with 3 μm ovalbumin-coated polystyrene beads (OVA-beads) for 25 min at 16°C, followed by 5 min at 37°C (Fig. 6A) and the non-internalised beads were then washed away. At different time points, we disrupted the cells to release phagosomes, stained them and analysed by flow cytometry. Neither the rate of Lamp1 acquisition nor that of ovalbumin degradation was affected by knocking out *Mpeg1* suggesting perforin-2 does not regulate phagosome maturation (fig. S6B and C). Using the perforin-2 αC-term antibody (Fig. 6B), we demonstrated that perforin-2 was rapidly recruited to phagosomes reaching its highest levels within 30 min after phagosome formation. This rapid acquisition suggests that perforin-2 is recruited prior to phagosome-lysosome fusion (*42*). Indeed, we could inhibit perforin-2 recruitment with an inhibitor of ARF GTPases Brefeldin A (BFA), which blocks protein trafficking through the early secretory pathway, suggesting perforin-2 is delivered to phagosomes from the Golgi. In phagoFACS experiments with the αMACPF Αb, perforin-2 acquisition was also BFA-dependent (Fig 6B and C). Unlike with the αC-term Ab, the αMACPF staining was restricted to mature, Lamp1+ phagosomes and increased over time while the signal for the cytosolic tail was gradually lost (Fig. 6C). Taken together, these data indicate that the C-terminal part of perforin-2 is cleaved off and degraded during phagosome maturation, while the ectodomain undergoes a conformational change in mature phagosomes, which exposes an epitope recognised by the αMACPF antibody.

**Fig 6.**
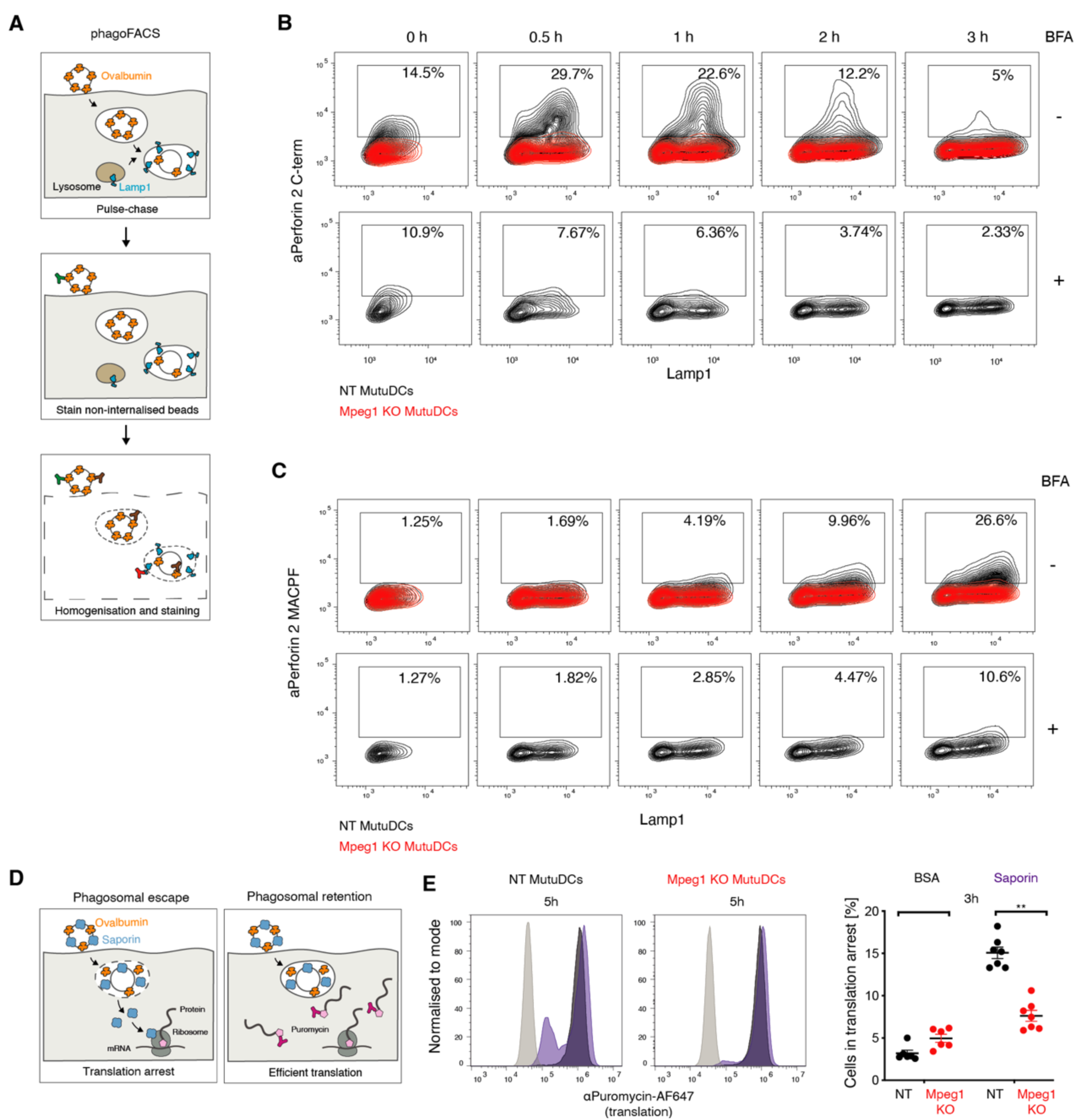
Perforin-2 undergoes pH-dependent proteolytic cleavage and mediates endocytic escape from antigen-containing phagosomes. **(A)** Schematic representation of the phagoFACS assay. Cells are pulsed with Ova-beads and allowed to internalise them. After an indicated chase period, non-internalised beads are marked with an αOvalbumin antibody. Following cell homogenisation, phagosomes are stained with antibodies against ovalbumin coupled to an alternative fluorophore and phagosomal markers. **(B and C)** *Mpeg1*^KO^ and NT MutuDCs were pulsed with Ova-beads and chased for the indicate time in the presence or absence of BFA. Isolated phagosomes were stained with antibodies against Lamp-1 and either the perforin-2 (B) C-terminus or (C) MACPF domain. Data are representative of three independent experiments. For gating strategy see fig. S6A. **(D)** Schematic representation of the saporin-bead puromycin assay. Cells are pulsed with saporin-conjugated Ova-beads and allowed to internalise them for the indicated time. Translation is then monitored with a 30 min puromycin chase. Incorporated puromycin can then be detected with an αPuromycin antibody and flow cytometry. **(E)** *Mpeg1*^KO^ and NT MutuDCs were pulsed for 3 or 5 h with saporin-conjugated Ova-beads and translation was monitored by a 30 min puromycin chase. Histograms are representative of two independent experiments each with two technical replicates. The quantification of translation inhibition represents three independent experiments each with at least two technical replicates, ns, not significant; *P<0.5; **P<0.01; ***P<0.001; ****P<0.0001 using an unpaired t test. For gating strategy see fig. S6D.

Finally, to confirm that perforin-2 contributes to the escape of phagocytic cargo, we conjugated saporin or BSA via thiol-ester bonds to OVA-beads and used them in the saporin-puromycin assay (Fig. 6D). We pulsed WT and *Mpeg1*^KO^ MutuDC with the beads and incubated them at 37°C for 5 hours. In WT, but not in *Mpeg1*^KO^ MutuDCs, saporin was able to escape from phagosomes leading to translation arrest (Fig. 6E).

### Perforin-2 is expressed in antigen-presenting cells and facilitates cross-presentation

To test whether the perforin-2-mediated endocytic escape pathway is involved in the delivery of antigens for cross-presentation, we generated knockout mice in which we disrupted the *Mpeg1* gene using a CRISPR-based approach (fig. S7A and B). Notably, knocking out *Mpeg1* did not result in any obvious disease phenotype. Furthermore, *Mpeg1*^-/-^ mice had immune cell frequencies comparable to WT mice (ffigig. S7C and D). Using intracellular staining, we confirmed that perforin-2 is expressed in spleen-resident cDC1s (Lin^-^CD11c^+^CX3CR1^-^XCR1^+^) but not in spleen-resident cDC2s (Lin^-^CD11c^+^CX3CR1^-^CD172a^+^), T cells, B cells or NK cells (fig. S8A). Interestingly, we also detected perforin-2 in other cell types, previously shown to cross-present *in vivo*, including splenic macropahges, plasmacytoid DC (pDCs), and a subpopulation of migratory cDC2s (Lin^-^CD11c^+^CX3CR1^+^CD172a^+^) (*43*–*45*) (Fig. 7A and fig. S8A). Indeed, splenic macrophages and pDCs, efficiently imported saporin into the cytosol *ex vivo* (fig. S8B and C). To confirm that loss of perforin-2 results in defective endocytic escape in primary cells, we performed the saporin-puromycin assay in bone marrow-derived DCs generated from WT and Mpeg1^-/-^ mice using Flt3-L and GM-CSF (Fig. 7B). In the BMDC^Flt3L/GM-CSF^ cultures, both cDC1s (CD11c^+^ XCR1^+^) and a large proportion of cDC2s (CD11c^+^ CD172a^+^) express perforin-2 (Fig. 7B). Saporin import into the cytosol is perforin-2-dependent in both subsets, as indicated by the differences in puromycin incorporation (Fig. 7C).

**Fig 7.**
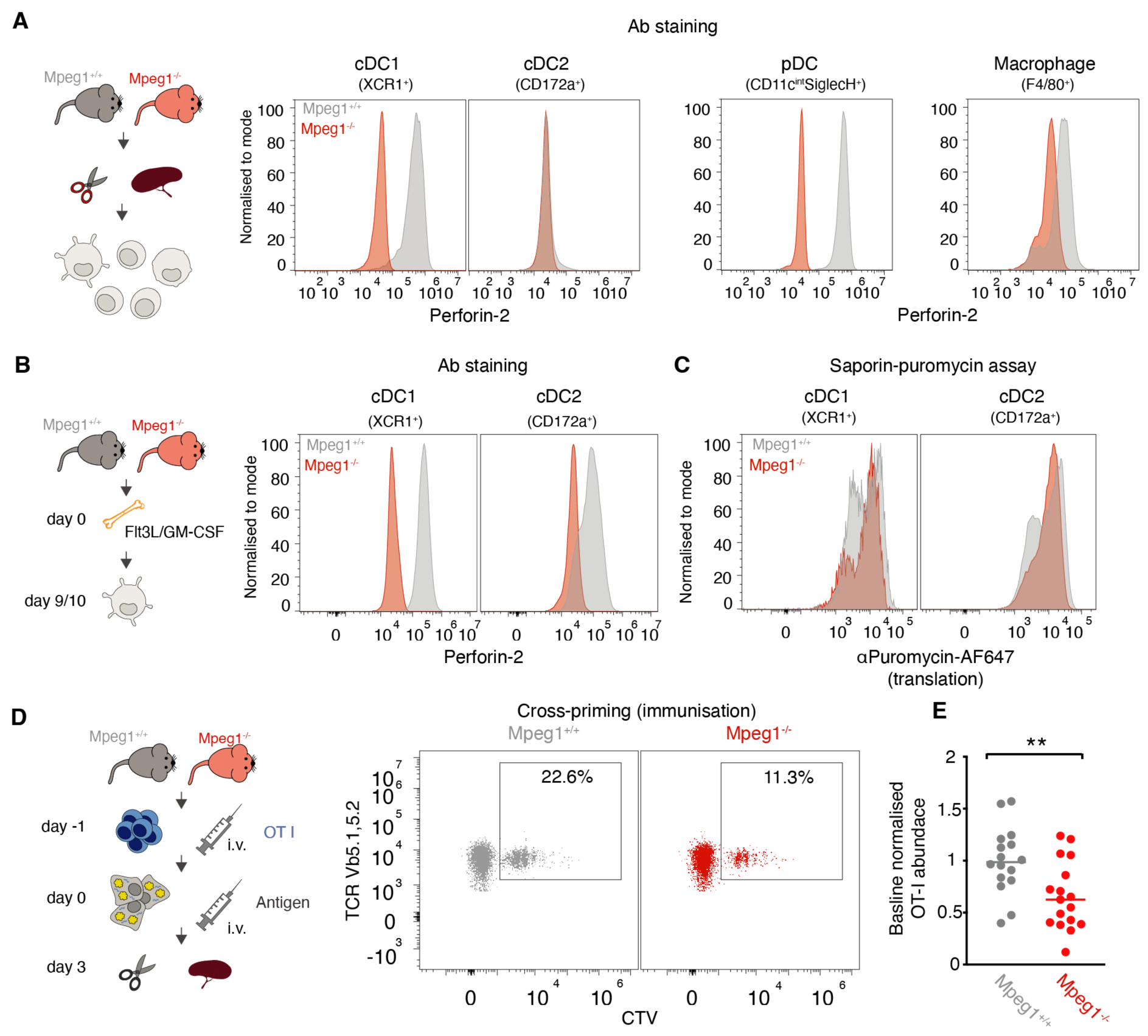
Antigen cross-presentation is impaired in vivo in the absence of perforin-2. **(A and B)** Perforin-2 expression was assessed by intracellular staining with an αPerforin-2 antibody and flow cytometry in (A) *Mpeg1^+/+^* and *Mpeg1*^-/-^ splenocytes and (B) Flt3-L/GM-CSF bone marrow cultures. In (A), cDC1s are defined as Lineage (CD3, CD19, NK1.1)^-^, F4/80^-^, CD11C^+^CX3CR1^-^, XCR1^+^, cDC2s as Lineage (CD3, CD19, NK1.1)^-^, F4/80^-^, CD11C^+^CX3CR1^-^, CD172a^+^, pDCs as Lineage (CD3, CD19, NK1.1)^-^, F4/80^-^, CD11c^int^SiglecH^+^ and macrophages as Lineage (CD3, CD19, NK1.1)^-^, F4/80^+^. In (B), cDC1s are defined as F4/80^-^, CD11c^+^XCR1^+^ and cDC2s as F4/80^-^, CD11c^+^CD172a^+^. For gating strategies see fig. S7C and S8D. **(C)** Flt3-L/GM-CSF bone marrow cultures from wild-type and *Mpeg1*^-/-^ mice were pulsed with saporin for 2 h, and translation was monitored by a 30 min puromycin chase. Flt3-L/GM-CSF cDC1s and cDC2s are defined as in (B, see also fig. S7C). Histograms are representative for three independent experiments. **(D)** Wild-type and *Mpeg1*^-/-^ mice were intravenously (i.v.) injected with 0.5x10^6^ CTV-labelled magnetically purified OT-I cells. One day later mice were i.v. injected with 1x10^6^ UVC-irradiated (240mJ/cm^2^) 3T3 cells, coated with 10 mg/mL ovalbumin and 0.5 mg/mL Poly(I:C). Three days later, OT-I proliferation was assessed by flow cytometry. OT-I are defined as Lineage (CD19, F4/80, CD11c)^-^, CD3^+^, CD4^-^CD8^+^, TCRvβ5.1, 5.2^+^TCRvα2^+^, CTV^+^. For gating strategy see fig. S8F. **(E)** Normalised OT-I counts three days after i.v. antigen injection. Each dot corresponds to an individual mouse, with three to five mice per group. For each experiment, OT-I counts per1x10^6^ splenocytes were normalised to the average of wild-type controls. Data from five independent experiments, ns, not significant; **P<0.01 using an unpaired t-test.

Finally, we asked whether perforin-2 mediated endocytic escape delivers antigens for cross-presentation. To assess the efficiency of antigen presentation *in vivo*, we adoptively transferred OT-I T cells expressing a TCR specific to ovalbumin peptide 257-264 displayed on H2Kb molecules and monitored OT-I proliferation following immunisation. Since cross-presentation of soluble ovalbumin, which can be processed by cell types other than cDC1s (*44*, *46*), was not significantly affected in the Mpeg1-/- mice (fig. S8F), we used dead cells as an antigen source. As anticipated, Mpeg1-/- had fewer OT-I T cells following immunisation with UVC-irradiated 3T3 fibroblasts coated with ovalbumin compared to WT mice. Thus, loss of perforin-2 leads to a defect in cross-presentation of cell associated antigens in vivo (Fig. 7D and E), suggesting that in cross-presenting cells, endocytic pores provide a route for cytosolic entry of antigens.

## Conclusions

In this study, we uncover a mechanism of endocytic escape that is governed by a cell type-specific pore-forming effector protein, perforin-2. The role of perforin-2 in the delivery of antigens for cross-presentation suggests that the immune system evolved to employ two related pore-forming effectors, perforin-1 and perforin-2, during different stages of adaptive immune responses. Perforin-1, expressed by cytotoxic T cells, is a well characterised effector employed for the delivery of granzymes into the cytosol of target cells (*47*). Our data, suggest that perforin-2 delivers endocytic contents into the cytosol of cross-presenting DCs, to enable the generation of MHC-I:peptide complexes for stimulation of naïve T cells.

We do not exclude the possibility that in different contexts, membrane destabilisation may also act to effectively deliver antigens for cross-presentation (*48*). Membrane integrity is regulated by a wide range of proteins involved in cell homeostasis (*49*, *50*), and as a result, different perturbations can result in leakage of endosomal contents into the cytosol under pathological conditions. Nevertheless, considering the restricted expression of *Mpeg1* to subsets of professional antigen presenting cells, our data points to at an important role of perforin-2 during the initiation of immune responses. The ability to genetically manipulate perforin-2-mediated endocytic escape also provides a new tool to explore the contribution of the escape pathway to anti-cancer and anti-viral immune responses *in vivo*.

## Acknowledgements

We thank members of the Kozik laboratory for discussions and feedback regarding this work; we also thank G.M. Griffiths and L.C. James for reading of the manuscript and their feedback. We thank N. Hacohen and T. Eisenhauer for their help setting up CRISPR-Cas9 screens. We thank the animal facility/ARES staff, genotyping facility, flow cytometry core and microscopy core at the MRC Laboratory of Molecular Biology for their technical assistance.

## Author contributions

PRS designed and performed experiments and wrote the manuscript. ML, designed and performed experiments and contributed to the manuscript. AD, JS and GHB carried out mass spectrometry and spatial proteomics analysis. BJT and WAM carried out Tau-HiBit assay. PAK performed experiments and provided scientific insight. PK supervised the study, designed, and performed experiments and wrote the manuscript.

## Additional Information

Correspondence and requests for materials should be addressed to Patrycja Kozik.

## Data and materials availability

All data are available in the main text or the supplementary materials.

## List of Supplementary Materials

**Materials and Methods**

**Figs. S1 to S8**

**Table S1**

**Table S2**

**Table S3**

**Table S4**

**Table S5**

**References only cited in Supplementary Materials**

## Materials and Methods

### Reagents

All reagents used in this study are listed in table S5. All antibodies used in this study are listed in the table S3.

### Cell Culture

MutuDCs were maintained in IMDM + Glutamax, 8% heat inactivated FSC, 10 mM HEPES, 500 μM β-mercaptoethanol, +/- penicillin (100 units/mL) / streptomycin (100 μg/mL). 3T3s, HEK293Ts and HeLas were maintained in DMEM + 10% FCS. Isolated primary T cells and bone marrow cells were maintained in RPMI-1640, 10% heat inactivated FSC, 10 mM HEPES, 500 μM β-mercaptoethanol, sodium pyruvate non-essential amino acids and penicillin (100 units/mL) / streptomycin (100 μg/mL).

For BMDC cultures, BM cells were resuspended in media supplemented with 5 ng/mL GM-CSF and 20 ng/mL Flt3-L. 15 x 10^6^ cells were seeded in a non-tissue culture treated 10 cm plate. Cells were differentiated for 9/10 days with the addition of 10 mL of fresh media with 5 ng/mL GM-CSF and 20 ng/mL Flt3-L on the fifth or sixth day and 5 mL of media containing 20 ng/mL GM-CSF added on the seventh day.

### Saporin assay

3 x 10^5^ cells were seeded per well in a treated 96-well U-bottom plate. Cells were pulsed for 2 h at 37°C with saporin. Cells were spun at 380g for 3 min and resuspended in DC media containing 0.01 mg/mL puromycin and incubated for 30 min at 37°C.

For the CRISPR/Cas9 screen, 3 x 10^6^ cells were seeded per well in a tissue culture treated 6-well plate. Cells were pulsed with 0.5 mg/mL saporin for 2 h at 37°C. Cells were then washed twice with media and incubated with puromycin at 0.01 mg/mL in DC media for 30 min at 37°C.

Saporin was labelled with an ATTO 550 protein labelling kit (Merck, 51146-1KT).

### Bead saporin

#### Bead preparation

Saporin or BSA were conjugated to Ova-beads through disulfide bonds. For efficient conjugation, free sulfhydryl groups were introduced by reacting 0.6 mL of 2.5 mg/mL saporin or BSA with 12 μL of 2 mg/mL Traut’s reagent in PBS + 2 mM EDTA for 60 min at room temperature. Excess reagent was removed using a Zeba spin desalting column equilibrated with PBS + 2 mM EDTA.

Ova-beads were resuspended in PBS + 2 mM EDTA and reacted with 1 mM SPDP for 30 min at room temperature. The beads were then washed twice in PBS + 2 mM EDTA and incubated with 1 mg/mL cysteine-modified saporin or BSA overnight at room temperature. After incubation, beads were washed twice in PBS and immediately used for phagocytosis.

#### Assay

The assay was performed in cell culture medium lacking β-mercaptoethanol. MutuDCs were collected, washed once in PBS and seeded in a 96-well U-bottom plate with 5 x 10^5^ cells per well in 100 µL cell culture medium. Saporin/Ova- or BSA/Ova-beads were diluted in cell culture medium such that adding 50 µL of each dilution to cells gave a 10:1 ratio of beads:cells. At the end of each incubation, puromycin was added at 0.01 mg/mL for 30 min and cells were washed in ice-cold PBS. Non-internalised beads were labelled in PBS containing 1% (vol/vol) BSA with a rabbit αOvalbumin antibody for 30min on ice followed by donkey αRabbit-AF555 for 30min on ice. After labelling dead cells with a fixable viability stain for 10 min on ice, cells were fixed and permeabilised using the BD Fix/Perm and Perm/Wash buffers. Puromycin incorporation was determined by staining with an αPuromycin-AF647 antibody in Perm/Wash buffer for 45 min on ice. Cells were analysed by flow cytometry.

### β-lactamase assay

4 x 10^6^ HeLa cells were pulsed with 800 uL of the CCF4 solution (prepared as in (*51*))) for 45 min, and washed by adding 5 ml PBS and spinning at 450 g, 15°C for 5 min. The cells were there resuspended in warm HeLa media with Probenecid at a 1:100 dilution (ThermoFished, P36400). To control for the background conversion of CCF4 observed in the absence of β- lactamase, each sample was split into two, media-only or media with β-lactamase (final concentration 2 mg/mL; P0389, Sigma). At each time point, 100 uL of the cell suspension was transferred into 100 uL of ice-cold PBS to stop trafficking. Finally, the cells were centrifuged for 2 min at 800 g, 4°C, stained with eFluor 780 live/dead solution containing Probenecid for 10 min on ice, centrifuged again, and resuspended in FACS buffer with Probenecid for flow cytometry.

### MutuDC survival assays

For saporin, gelonin and cycloheximide treatments, 5 x 10^4^ cells were seeded per well in a treated 96-well U-bottom plate. Cells were incubated with the different treatments for 24 hours. Cells were washed twice in PBS and then stained with a fixable viability stain. Cell viability was assessed by flow cytometry.

For treatment with Poly (I:C) 2.5 x 10^4^ cells were seeded per well in a treated 96-well U-bottom plate. Poly (I:C) was added and cells were incubated for 48 hours. Cells were washed in twice in PBS and then stained with a fixable viability stain. Cell viability was assessed by flow cytometry.

For treatment with Bleomycin A1, 2 x 10^5^ cells were seeded per well in a treated 96-well U-bottom plate. Bleomycin A1 was added and cells were incubated for 48 hours in the Incucyte®. The increase in cell area covered by the cells was monitored for each well and normalised to untreated cells.

### PhagoFACS

#### Bead preparation

To generate Ova-beads, amino-modified microspheres with a diameter of 3-μm were washed twice in PBS and preactivated with 8% (vol/vol) glutaraldehyde for 4 h at room temperature. Preactivated beads were washed once in PBS and then incubated overnight at 4°C with ovalbumin at a concentration of 0.5 mg/mL in PBS. After incubation, beads were quenched in 0.4 M glycine in PBS, washed twice in PBS and used immediately for phagocytosis.

#### PhagoFACS assay

MutuDCs were collected, washed once in PBS and resuspended in ice-cold internalisation medium (CO2-independent medium containing 1X GlutaMAX) to a density of 20 x 10^6^ cells/mL. Ova-beads were added at a 10:1 ratio of beads:cells and incubated for 25 min at 16°C followed by a 5min incubation at 37°C to allow phagocytic binding and internalisation of beads. To remove non-internalised beads, cells were first washed twice with 10 mL ice-cold PBS at 100 g for 4 min at 4°C and then resuspended in 1 mL PBS, applied to a 5 mL FCS cushion and centrifuged at 150 g for 4 min at 4°C. The cell pellet was then resuspended to 20 x 10^6^ cells/mL in cell culture medium (containing 5 µM BFA for drug treated cells) and divided into different time points comprising 5 x 10^6^ cells each. The chase was performed at 37°C for different periods of time and stopped by adding ice-cold PBS. Non-internalised beads were labelled by staining with a goat αOvalbumin antibody for 30min on ice followed by an αGoat-AF488 antibody for 30min on ice. Cells from each time point were resuspended in 0.5 mL homogenization buffer (250 mM sucrose, 3 mM imidazole, 2 mM DTT, 2 mM PMSF and 1X protease inhibitor cocktail, ph 7.4) and passed 25 times through a 22-G needle. Intact cells and debris were pelleted by centrifugation at 150 g for 4min and the phagosome-containing post-nuclear supernatants transferred to a V-bottom 96-well plate. The enriched phagosomes were washed with PBS containing 1% (vol/vol) BSA and stained with different primary antibodies overnight at 4 °C. The next day, the samples were incubated with appropriate secondary antibodies for 45 min on ice. Phagosomes were analysed by flow cytometry.

### Tau entry assay

Tau entry assays were performed as previously described (*34*). Briefly, HEK 239T cells expressing NLS-eGFP-LgBiT (NGL) were transduced with lentivirus harbouring human*Mpeg1*^IRES^BFP or BFP only under the control of an SFFV promoter. Approximately 16 h prior to assay, 2x10^4^ cells were seeded into a white 96-well plate coated with poly-L-lysine. The media was replaced 16 h later with recombinant 0N4R-P301S-tau-HiBiT (tau-HiBiT) protein at desired concentration in Assay Medium composed of CO2 independent medium supplemented with 1% penicillin-streptomycin, 1 mM GlutaMAX and 1 mM sodium pyruvate. Cells were washed once with PBS, and incubated in substrate solution (Assay Medium, LCS buffer and Live Cell Substrate) at RT for 5 min before loading onto a pre-warmed clarioSTAR Microplate Reader at 37°C. The plate was mixed by 200 RPM double orbital shaking for 10 seconds prior to signal acquisition by spiral average (NanoLuc setting, 470 nM). Post-signal acquisition, cell viability per well was acquired by incubation with PrestoBlue Viability Reagent according to manufacturer instructions. The plate was then loaded onto the ClarioSTAR Microplate Reader and fluorescence read by excitation wavelength of 560 nm and emission at 590 nm.

### 3T3 UVC-irradiation and antigen coating

1.5 x 10^6^ 3T3 cells were plated in 10 cm plate 16 hours prior to irradiation. For irradiation, media was removed and 5 mL PBS was added. Cells were UVC irradiated (240 mJ/cm^2^) with a UVP Crosslinker (AnalytikJena). Media was then replenished, and cells were incubated for 16 h at 37°C.

For antigen coating, the supernatant, the PBS wash and the trypsinised UVC-irradiated 3T3s were collected and resuspended at 10 x 10^6^ cells/mL in 10mg/mL ovalbumin and 0.25 mg/mL Poly(I:C). Cells were incubated for 1 h at 37°C. Cells were washed three times in ice-cold PBS and resuspended for injections.

## OT-I CTV labelling

Staining was performed with a CellTrace Violet Cell Proliferation Kit. Isolated cells were resuspended at 5 x 10^5^/mL in a 2.5 μM CellTrace Violet solution, and incubated at 37°C for 20 min in the dark. 10% v/v FSC was added, and cells were incubated a further 5 min at 37°C. Cells were centrifuged at 300 g for 5 min, resuspended in T cell media and incubated at 37°C for 10 min.

## Imaging

### Mpeg1 trafficking

1.5 x 10^5^ MutuDCs were plated on poly-L-lysine coated μ-slide 8 well dish and allowed to adhere at 37°C overnight. To label acidic compartments, the cell culture medium was replaced with fresh medium containing 1 µM LysoTracker Red and cells were incubated for 30 min at 37°C. Cells were then washed twice with PBS and fixed in 4% formaldehyde for 10 min at RT. After two washes with PBS, the samples were permeabilised with 0.15% Triton X-100 in PBS for 10 min at RT followed by three washes with PBS. To block unspecific binding and quench excess formaldehyde, the cells were incubated in blocking buffer (1% BSA, 0.3M glycine, 0.1% Tween 20 in PBS) for 30 min. Primary and secondary antibody incubations were performed in PBS containing 1% BSA for 1 hr at room temperature with three washes in PBS after each incubation. Nuclei were labelled by incubating the cells for 10 min with DAPI followed by two washes in PBS. Images were acquired on a VisiTech iSIM swept field confocal super resolution system coupled to a Nikon Ti2 inverted microscope stand equipped with a 100x/1.49 NA SR Apo TIRF objective lens. The images were processed and analysed in Fiji.

### Galectin 3 endosomal recruitment

1 x 10^5^ MutuDCs were plated on a μ-slide 8 well dish and allowed to adhere at 37°C overnight before treating them with 33 μM GPN for 10 min at 37°C. Cells were washed three times in PBS and fixed in 4% paraformaldehyde for 10 min at RT. Paraformaldehyde was washed away before permeating cells with 0.1% Triton-X100 for 10 min at RT. Cells were washed 3 times in PBS, and incubated for 30 min at RT with blocking buffer (1% BSA, 0.3M glycine, 0.1% Tween 20 in PBS). After three PBS washes, cells were stained with αGalectin3 for 40 min and then washed 3 times in PBS. Cells were then stained with donkey αMouse-Af647 and then washed three times in PBS. Images were acquired on a Zeiss 780 inverted confocal microscope. The images were processed and analysed in Fiji.

## Mice

All mice were bred/maintained in pathogen-free conditions by the Medical Research Council ARES facility. Experiments were approved by the LMB Animal Welfare and Ethical Review Body and the UK Home Office. MRL/MpJ-Fas^lpr^/J (The Jackson Laboratory, 000485) mice were a gift from L.C James (MRC Laboratory of Molecular Biology). C57BL/6, MRL/MpJ-Fas^lpr^/J, OT-I and *Mpeg1*^-/-^ mice used were 8 to 12 weeks old at the time of the experiment.

## Mouse tissues

Spleens were perfused with a solution of RPMI-1640, 0.1 mg/mL Liberase-TL and 0.1 mg/mL DNAse I, minced and digested for 25 min at 37 °C. Heat-inactivated FCS was added (10% v/v) to stop the digestion, before filtering tissues through a 70 μM filter. Red blood cells were then lysed with red blood cell lysis buffer hybrid-max. To obtain the bone marrow, femurs and tibias from mice were cut at both ends, and the bone marrow was flushed into BMDC media by brief centrifugation at 10,000g. Red blood cells were lysed with red blood cell lysis buffer hybrid-max. Tissue CD11c+ cells were enriched using a Pan-Dendritic Cell Isolation Kit. OT-I T cells were obtained from OT-I spleens and lymph nodes. Both tissues were mashed and filtered through a 70 μM filter. OT-I T cells were enriched using a EasySep Mouse Naïve CD8+ T Cell Isolation Kit.

## In vivo immunisation

8–12 week old male and female mice were i.v. injected 0.5 x 10^6^ OT-I cells. The following day mice were injected with either 100 μg of ovalbumin + 50 μg Poly(I:C), or with 1x10^6^ UVC-irradiated 3T3s coated as described with 10 mg/mL ovalbumin and 0.5 mg/mL Poly(I:C). Three days later spleens were isolated and OT-I T cell abundance was assessed by flow cytometry.

## CRISPR/Cas9 knockout mice generation

Both the crRNAs and the tracrRNA (TRACRRNA05M-5NMOL) were, ordered from Sigma-Aldrich, (HPLC-purified and 2-O-Methyl capped).

The sequence of the crRNA (functional) was GCUCAGCUUGGGGUUUACGA. Due to a design error, an additional non-functional crRNA (CGAUGAAGUGUAUACUAUUC) was injected. This crRNA has no targets in the mouse genome according to the Wellcome Sanger Institute Genome Editing tool (https://wge.stemcell.sanger.ac.uk/).

All oligos and Cas9 nuclease were initially resuspended in RNAse free 10 mM TrisHCL, 0.1 mM RNAse-free EDTA at 500 ng/mL and 200 ng/mL, respectively. Oligos and Cas9 were mixed in RNAse free 10 mM TrisHCL, 0.1 mM EDTA and incubated at room temperature for 15 min (final concentrations 20 ng/mL for both oligos and Cas9). The Cas9-gRNA complexes were injected into a C57/Ola zygote, which was then allowed to recover for 3 h prior to its transfer into a CD1 surrogate.

We assessed editing efficiency by Sanger sequencing of a 560 bp amplified DNA sequence from the target region (primers can be found in Supplementary Table 1). In doing so we identified three recurrent mutations in the different founder mice, which consisted of a 40-base deletion affecting CDS bases 526 to 566, a 17-base deletion spanning CDS bases 525 to 541, and lastly a 16-base deletion of CDS bases 524 to 540. Mice with the 17-base deletion were used for experiments.

## Plasmids

Plasmids used for generation of the constructs below are listed in the table S4.

### mMpeg1 Overexpression Construct

Mouse *Mpeg1* sgRNA-resistant DNA, was generated using GeneArt Gene Synthesis (sequence in table S4). Appropriate homology arms were added to *Mpeg1* cDNA by PCR with KOD Xtreme Hot Start DNA (primer sequences in table S4). *Mpeg1* DNA was inserted by Gibson assembly into pHR-scFv-GCN4-sfGFP-GB1-NLS-dWPRE.

### mScarlet only construct

pHR-scFv-GCN4-sfGFP-GB1-NLS-dWPRE was digested with MluI-HF and NotI-HFat 37°C °for 1 hour. mScarlet, with appropriate homolgy arms (primer sequences in table S4) was added by Gibson assembly.

### Mpeg1^IRES^-mScarlet construct

pHR-scFv-GCN4-sfGFP-GB1-NLS-dWPRE was digested with MluI-HF and NotI-HF at 37°C

°for 1 hour. For the *Mpeg1*^IRES^mScarlet, the IRES promoter and mScarlet were added by Gibson assembly to the *m*Mpeg1 overexpression construct. The primers used to add the homology arms can be found in table S4.

### BFP only construct

For the generation of the BFP-only construct, mTagBFP2 (table S4) was cloned in replacing mScarlet in the mScarlet only construct using MluI and NotI sites.

### humanMpeg1^IRES^-BFP construct

This plasmid was derived from the *Mpeg1^I^*^RES^-mScarlet by replacing murine Mpeg1 for a human Mpeg1 IDT gene block (table S4) using KpnI and NotI sites. mScarlet was then replaced with ^IRES^mTagBFP2 using Mlul and NotI sites (table S4).

### WT-mMpeg1restr^IRES^-mScarlet and mMpeg1G212V/A213V^IRES^-mScarlet and mMpeg1K251C/G286C^IRES^-mScarlet constructs

For this construct wildtype murine *Mpeg1* was replaced with a Twist gene fragment encoding wild-type murine *Mpeg1* with a silent point mutation at codon positions 495 to introduce a SbfI site (denoted mMPEG1restr).

mMPEG1^G212V/A213V-IRES^-mScarlet and mMPEG1^K251C/G286C-IRES^-mScarlet were generated by cloning IDT gBlocks harbouring the respective mutations into mMPEG1restr^IRES^-mScarlet using the BstXI and SbfI sites. Sequence for these Twist gene fragments can be found in table S4.

### Mpeg1 sgRNA construct

The lentiBFP plasmid (a gift from A.N.J McKenzie, MRC Laboratory of Molecular Biology) was digested with BsmBI and dephosphorylated with CIP. Phosphorylated and hybridised oligos, corresponding to the sgRNA sequences, were ligated into the plasmid using T4 DNA ligase.

## Lentiviral Production

3x10^6^ HEK293T cells were seeded per 10 cm dish. After 24 hours, cells were transfected with the sgRNA plasmid and lentiviral packing plasmids (psPAX2 and VSVG-Pmd2) using TransIT-LT1 transfection reagent. Media was changed 18 hours post-transfection to DMEM, 10% FCS, 1% BSA. Virus-containing supernatant was first collected 48 h post-transfection and media was replenished. This first virus batch was kept overnight at 4 °C. A second virus batch was collected the following day (72 h post-transfection), and both aliquots were mixed. Cellular debris was removed by centrifugation at 380 g for 5 min, before concentrating the virus by centrifugation at 3000 g for 45 min in Amicon Ultra-15 centrifugal filters.

## CRISPR/Cas9 Screen

### sgRNA library design

Microarray expression data for splenic cDC1s (CD8α^+^ DCs) and cDC2s (CD4^+^ DCs) was downloaded from Immgen (www.Immgen.org) (*35*) on Feb 5^th^ 2017. Genes with log2(cDC1/cDC2) expression ratio > 1.3 were selected for the minilibrary (table S1). Four sgRNA sequences per gene were picked from the genome-wide Brie library (*36*). For genes not present in the Brie library, the sgRNAs were designed using the Broad Institute Genetic

Perturbation Platform portal: https://portals.broadinstitute.org/gpp/public/analysis-tools/sgrna-design. The resulting library targeted 281 genes (table S1).

### sgRNA library generation

The oligos to generate the sgRNA library were ordered from Twist Bioscience as part of a larger pool containing several libraries. The sequences are listed in table S1. The oligo pool was resuspended to 53 nM and minilib-PCR1 primers (table S1) were used to amplify the library.

The LentiBFP vector was digested with BsmBI, and a gibson reaction was set up with the library inserts. The Gibson reaction mix was electroporated into EnduraTM competent cells and electroporated at 1.8 kV/600Ο/10μF. Bacteria were allowed to recover in recovery media for 1 h at 37°C, before plating them in a Luria broth (LB) lennox NuncTm Square BioAssay dish and growing them overnight at 30°C. Bacteria were then collected by washing the plate 5 times with 5 mL LB. Bacteria were centrifuged at 4000 g for 15 min, before removing the supernatant and freezing the pellet. For DNA purification, the bacteria pellet was thawed, and DNA purified using a QIAGEN Plasmid Maxi Kit.

### Lentiviral Transduction for the CRISPR-Cas9 screen

Cas9 expressing MutuDCs were seeded at 2x10^6^ per 10 cm dish. Virus was added at a 0.3 MOI. Two days after transduction BFP positive cells were sorted and allowed to recover. Library representation was maintained at 800x.

### Preparation of libraries for next generation sequencing

Genomic DNA was isolated using a DNeasy Blood and Tissue Kit. Samples were digested for 4 hours with Proteinase K. sgRNAs were then amplified in a two-step PCR with Herculase II Fusion DNA Polymerase. For the first PCR a maximum of 5 ug of DNA per 50 μL reaction was amplified using all available genomic DNA and libgen-PCR1 primers (table S1). The reaction mixed consisted of 10 μL 5X Herculase II reaction buffer, 0.5 μL dNTP mix, 0.5 μL Herculase II fusion DNA polymerase and 1.25 mL of 10 μM both the forward and reverse primers. PCR cycling conditions were: 1 min at 95 °C; 18x (30 s at 95°C, 30 s at 55 °C, and 30 s 72 °C); 10 min at 72°C).

The number of cycles for the second PCR was determined by qPCR using a KAPA Library Quantification Kit. Cycling conditions were: 2 min at 95 °C, followed by 30 s at 95 °C, 30 s at 53 °C and 30 s at 72 °C for a total of 40 cycles. Samples with similar CT values were then pooled for the second PCR.

Following sgRNA amplification from gDNA, a second PCR was performed to barcode the sgRNAs using 10 μL from the first PCR. PCR cycling conditions were: 2 minutes at 95 °C, followed by 30 s 95 °C, 30 s at 53 °C and 30 s at 72 °C for the number of cycles determined by the previous qPCR step, and a final 10 min at 72 °C. The amplified products were separated using a 2% agarose gel and purified using a Gel Extraction Kit. Agarose was melted at 40 °C instead of the recommended 50 °C. Samples were then further purified using a Charge Switch PCR Clean-Up Kit. Finally, the concentration of DNA fragments containing P5 and P7 adaptors was determined via qPCR using KAPA Library Quantification Kit. This ensured adequate cluster density during NSG. This was done using a KAPA Library Quantification Kit. Samples were analysed by SE50 sequencing on a HiSeq4000 sequencer loaded with 15% spike-in PhiX Control Library (Illumina).

## Analysis of the CRISPR-Cas9 screening data

### Processing of the sequencing data

The analysis pipeline was adapted from (*52*). Demultiplexing was performed using the demuxFQ package. Next, the fastq files were processed using cutadapt-1.4.1 to remove the flanking sequences: GACGAAACACCG and GTTTTAGA on the 5’ and 3’ end respectively (analysis parameters: -e 0.2 --minimum-length 20 --discard-untrimmed). The trimmed 20 bp long reads were matched to the library of reference sgRNA sequences and counted (analysis parameters: -f -v 0 -m 1 --norc -a --best --strata --un). Raw counts for each sgRNA are provided in table S1.

### Hit calling

Each screen repeat was analysed initially using a modified stat.wilcox function from the caRpools (v 0.83) package with the following parameters and modifications. Guides with less than 20 counts in either of the populations selected for comparison were excluded from the analysis. The counts were then normalized to median of the population. The function returns enrichment score for each population of four sgRNAs targeting one gene relative to 100 random guides. The p-values are calculated using a two-sided Mann-Whitney-U test and non-adjusted P-values were used in the next step of the analysis.

To combine the data from the biological replicates of the screen, for each gene we calculated the mean enrichment score and used Fisher’s method to combine the p-values. P-values were then adjusted using the Benjamini-Hochberg method. Genes with enrichment scores greater than 0.5 and adj p-values < 0.01 were considered as hits.

## Recombinant perforin-2 cleavage

### Expression and purification of recombinant perforin-2

The murine perforin-2 ectodomain (amino acids 20-652), tagged with an N-terminal signal peptide and a C-terminal hexahistidine tag was introduced into vector pHL-sec. Expi293F cells at 2x10^6^/mL were transfected with 3.8 mg of plasmid in the presence of 10.2 mg of Polyethyleneimine Max. Three to four hours later, valporic acid was added at a final concentration of 3.5 mM. Cells were allowed to grow for a further 5 days. Culture media was collected, centrifuged for 2 h at 4000g and filtered through a 0.22 μM membrane. An ÄKTA flux^TM^ (Cytiva Life Sciences) was used for sample concentration (with a 10kDa cut-off filter) and buffer exchange to Buffer A (25mM Tris pH 7.5, 500Mm NaCL, 10 mM imidazole). The sample was then loaded onto a Ni-NTA column at RT. The column was washed, sequentially, with 10 mM and 40 mM imidazole, before collecting the perforin-2 elute with a 200 mM imidazole wash. The perforin-2 fractions were then dialyzed in PBS and concentrated to 2 mg/mL.

### Asparagine Endopeptidase cleavage reactions

AEP (specific activity 350 pmol/min/μg) was resuspended in activation buffer (0.1 M NaOAc, 0.1 M NaCl, pH 4.5) at 50 μg/mL. Prior to the cleavage reactions, ΑEP was incubated for 4 h at 37°C. For the *in vitro* cleavage reactions, AEP was diluted in assay buffer (50 mM MES, 250 mM NaCl, adjusted pH 5.5) and 35 or 175 μU were added to a 50 μL final reaction volume. To ensure AEP was active, cleavage of a fluorogenic substrate was confirmed using manufacturer’s protocols. For perforin-2 cleavage, 4 μL of purified perforin-2 was cleaved in the absence or presence of AEP inhibitor peptide at 0.5 mg/mL. Reactions were allowed to proceed for 2 h at 37°C before being terminated by addition of denaturing agent.

## Whole cell lysate proteomic analysis

### Cell lysis and in-solution digestion of proteins

Cell pellets were thawed, lysed in 200 µl 2.5% (w/v) SDS/50mM Tris pH 8 and incubated at 72°C for 5min. Lysates were then sonicated at 4 °C (three times 5s bursts with an amplitude of 10µm) to break-up DNA. Estimations of protein concentrations were made using a Pierce BCA Protein Assay Kit. For each sample, 100 µg protein was precipitated by the addition of 5 volumes of ice-cold acetone, incubated at −20°C overnight and pelleted by centrifugation at 4°C for 5min at 10,000×g. All subsequent steps were performed at room temperature. Precipitated protein pellets were air-dried for 5min, resuspended in 50 µl digestion buffer (50mM Tris pH 8.1, 8M Urea, 1mM DTT) and incubated for 20min. Protein was alkylated by addition of 5mM iodoacetamide for 20min (in the dark) and then enzymatically digested by addition of LysC (1mg per 50mg of protein) for an overnight incubation. Digests were then diluted four-fold with 50mM Tris pH 8.1 before addition of Trypsin (1mg per 50mg of protein) for 4 hours. The peptide mixtures were then acidified to 1% (v/v) TFA in preparation for peptide purification and fractionation.

### Peptide purification and fractionation

For each sample, 20 µg peptides were loaded onto an SDB-RPS StageTip for peptide cleanup and triple fractionation as previously described (*53*). StageTips were activated by washing with 100 µl acetonitrile, followed by 100 µl 30% (v/v) methanol, 1% (v/v) TFA, and then 100 µl 0.2% (v/v) TFA. Peptide mixtures in 1% TFA were loaded onto activated StageTips and washed with 100 µl isopropanol and then 100 µl 0.2% (v/v) TFA. Peptides were then eluted successively using 20µL SDB-RPSx1 (100mM ammonium formate, 40% (v/v) acetonitrile, 0.5% (v/v) formic acid), then 20µL SDB-RPSx2 (150mM ammonium formate, 60% (v/v) acetonitrile, 0.5% formic acid), then 30µL Buffer X (80% (v/v) acetonitrile, 5% (v/v) ammonium hydroxide). Peptides were dried in a centrifugal vacuum concentrator, resuspended in 10µL Buffer A* (0.1% (v/v) TFA, 2% (v/v) acetonitrile) and stored at −20°C until analysis by mass spectrometry.

### Mass spectrometry

500 ng of peptides were loaded on a 50 cm by 75 µm inner diameter column, packed in-house with 1.8 µm C18 particles (Dr Maisch GmbH, Germany). Peptide separation by reverse phase chromatography was performed using an EASY-nLC 1000 (Thermo Fisher Scientific), running a linear gradient over 95 min at 300 nl/min flow rate and 55 °C. The gradient ran from buffer A (0.1% (v/v) formic acid) containing 5% buffer B (80% (v/v) acetonitrile, 0.1% (v/v) formic acid) to buffer A containing 30% buffer B. Runs were separated by 5 min wash-outs with 95% buffer B and re-equilibration. The LC was coupled to a Q Exactive HF-X Hybrid Quadrupole-Orbitrap mass spectrometer via a nanoelectrospray source (Thermo Fisher Scientific). MS data were acquired using a data-dependent top-15 method that dynamically excludes precursors picked during the last 30 seconds. MS1 survey scans were acquired at a resolution of 60,000 in a 300-1650 Th range. The maximum injection time was 20 ms for up to 3e6 target ions, as determined with predictive automatic gain control. Sequencing was performed via higher energy collisional dissociation fragmentation of ions isolated from a 1.4 Th window. The maximum injection time was 28 ms for 1e5 target ions. MS2 fragment scans were acquired at a resolution of 15,000 in a 200-200,000 Th range. The minimum predicted ion count to be reached per injection was 2.9e3.

### Processing of mass spectrometry data

Mass spectrometry raw files were processed in MaxQuant Version 1.6.10.43 (*54*), using the human SwissProt canonical and isoform protein database, retrieved from UniProt (2019_11_29; www.uniprot.org). Label-free quantification was performed using the MaxLFQ algorithm(*55*). No matching between runs was used. To detect peptide fragments resulting from other cleavage events than the in-solution digest with LysC and trypsin, the enzyme mode was set to semi-specific. LFQ minimum ratio count was set to 1 and default parameters were used for all other settings.

To assess abundance of peptides independent of protein abundance changes, peptide intensities were divided by their corresponding protein intensities. Data were filtered for 3 valid values in at least one experimental condition (n=52160) and then subjected to a two-sided student’s t-test. Multiple hypothesis correction was done by permutation based FDR with s0=0.1 using Perseus (*56*)

### Flow cytometry

All flow cytometry staining was done on ice. For cell surface staining, nonspecific binding was blocked using αCD16/CD32. Cells were then stained with the corresponding antibodies for 30-60 min (table S3). Following antibody staining, cells were washed three times with PBS, 1mM EDTA, 1% FCS. For live/dead staining cells were either resuspended in a (0.1

μg/mL) DAPI PBS, 1mM EDTA, 1% FCS solution or stained for 10 min with a fixable viability dye (table S3). Intracellular staining was done using BD Cytofix/Cytoperm Fixation/Permeabilization Kit. Data was acquired on a LSRFortessa (BD) or a CytoFLEX (Beckman Coulter) and analysed on FlowJo 10 (BD). For cell sorting, cells were sorted into serum coated polypropylene tubes using a SY3200 Cell Sorter (Sony).

### Western Blot

Cell pellets were lysed in RIPA buffer with protease inhibitor for 20 min at 4°C while shaking at 800 rpm. Lysates were centrifuged at 20,000g for 10 min at 4°C, the supernatant was then transferred to a clean 1.5 mL Eppendorf. Lysates were mixed with LDS NuPAGE + Bolt reducing agent and incubated at 70°C for 10 min. Samples were loaded onto an SDS-Page gel and transferred onto a nitrocellulose membrane using an iBlot1 (Invitrogen). The membrane was blocked for 1 h with 5% milk solution in PBS, 0.01% Tween (PBST), and blotted with the primary antibody overnight at 4°C. Secondary staining was performed for 1h at RT. Chemiluminescence was detected with ECL Prime Western Blotting Detection Reagent.

## Supplementary Figures

**Fig S1.**
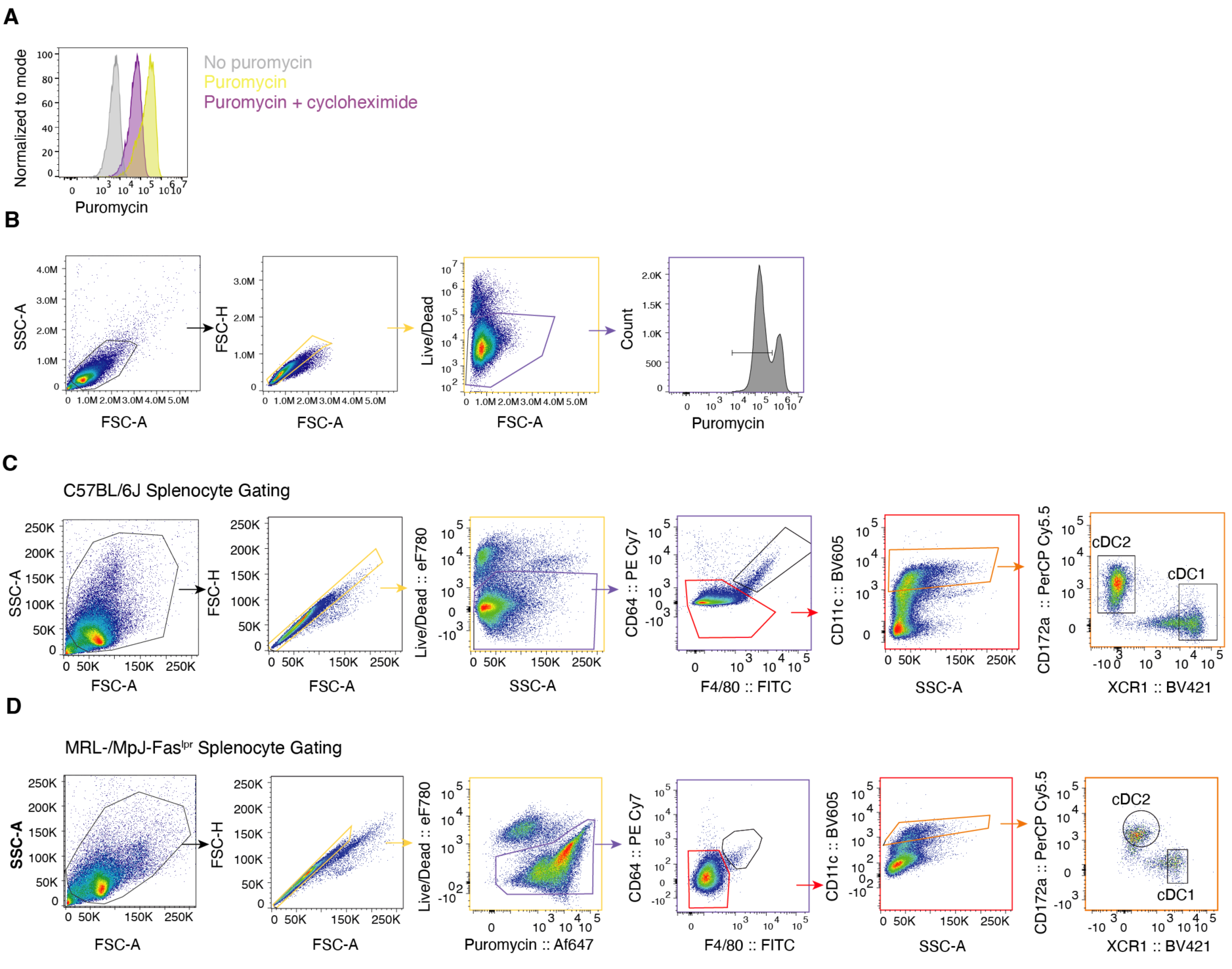
Monitoring puromycin incorporation allows to track changes in translation. **(A)** Monitoring puromycin incorporation by flow cytometry. MutuDCs were incubated for 2 h at 37°C with 10 μg/mL cycloheximide followed by a 30 min 0.01 mg/mL puromycin chase in the presence or absence of 10 μg/mL cycloheximide. Puromycin incorporation was monitored with a αPuromycin antibody. Histograms are representative for three independent experiments. **(B)** Flow cytometry gating strategy to monitor translation inhibition in MutuDCs. **(C)** Flow cytometry gating strategy for C57BL/6J splenic cDC1s and cDC2s for the *ex vivo* saporin assay. **(D)** Flow cytometry gating strategy for MRL/MpJ-Fas^lpr^ splenic cDC1s and cDC2s for the *ex vivo* saporin assay.

**Fig S2.**
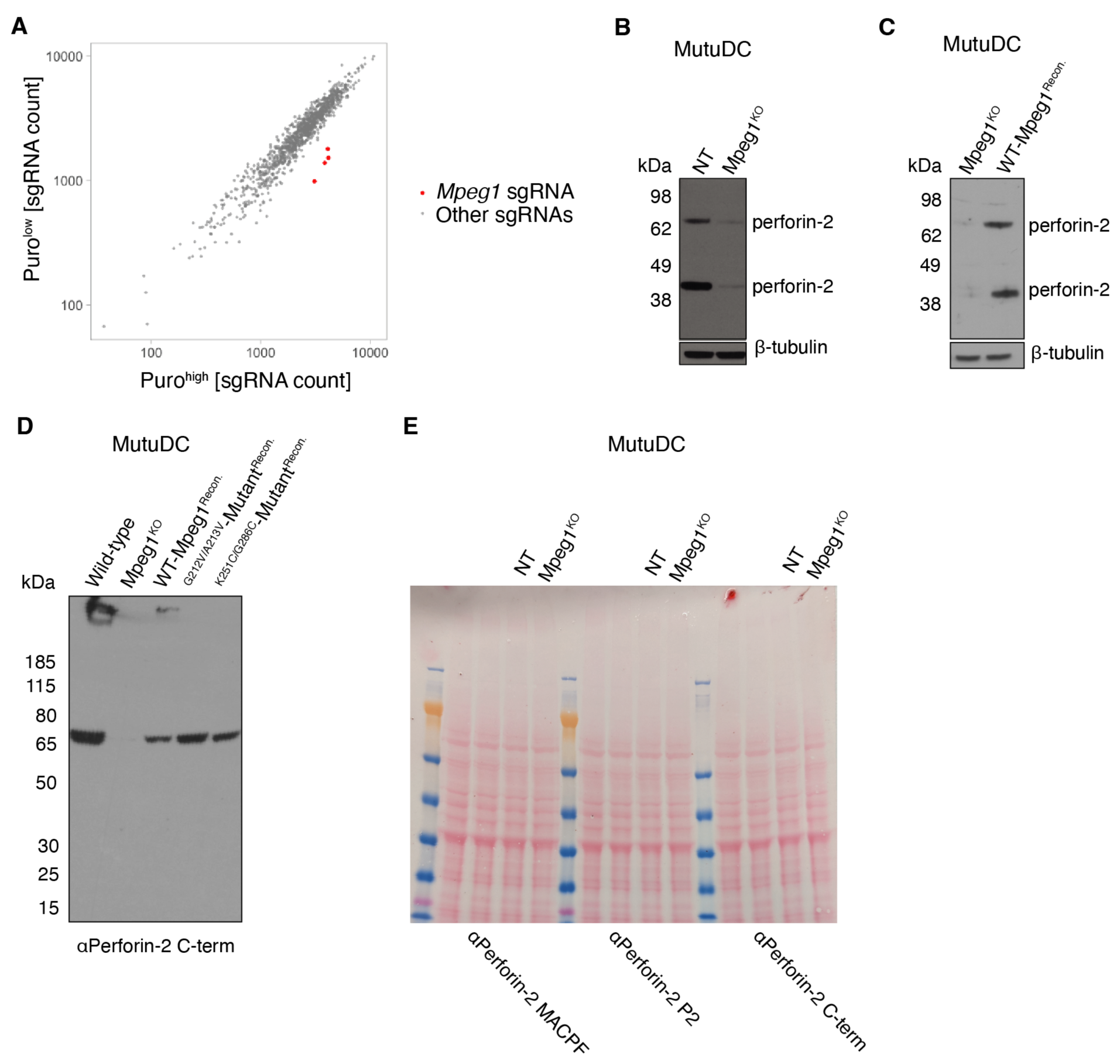
Identification of *Mpeg1* in the CRISPR-Cas9 genetic screen and characterization of the *Mpeg1* KO and rescue MutuDC lines. **(A)** Relative abundance of sgRNAs in puro^high^ and puro^low^ populations from the saporin-puromycin-based genetic screen. The screen was performed in three biological repeats. The sgRNA counts from each population in each screen replicate were normalised, and the average of the counts for puro^high^ and puro^low^ populations is plotted. Each dot corresponds to one sgRNA. The sgRNAs targeting *Mpeg1* are highlighted in red. **(A)** Perforin-2 protein levels in *Mpeg1*^KO^ and NT MutuDCs were assessed by Western blot under reducing conditions. β-tubulin was used as a loading control. **(B)** Perforin-2 protein levels in *Mpeg1*^KO^ and *Mpeg1*^KO^ MutuDCs complemented with sgRNA resistant *Mpeg1* were assessed by Western blot under reducing conditions. β-tubulin was used as a loading control. **(C)** Protein levels of WT and perforin-2 mutants in reconstituted *Mpeg1*^KO^ MutuDCs were assessed by Western blot under non-reducing conditions. **(D)** Ponceau staining corresponding to the Western blot in Fig. 5C.

**Fig S3.**
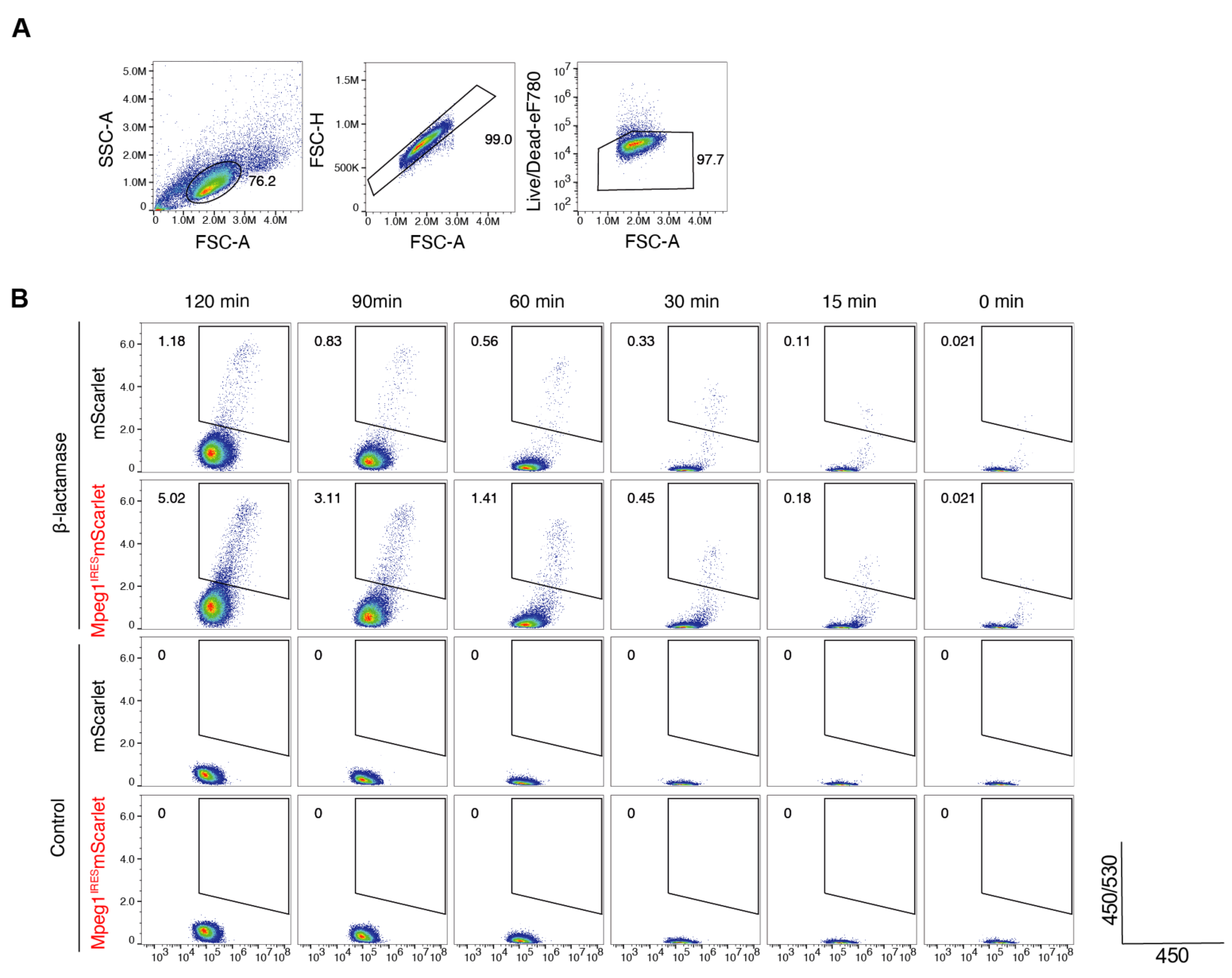
Gating strategy for the CCF4 β-lactamase assay. **(A)** Flow cytometry gating strategy for the CCF4 β-lactamase assay in mScarlet and Mpeg1^IRES^-mScarlet HeLa cells. **(B)** Representative gating for CCF4 cleavage for each condition. Gates were defined to exclude spontaneous conversion of CCF4 in the absence of β-lactamase over time.

**Fig S4.**
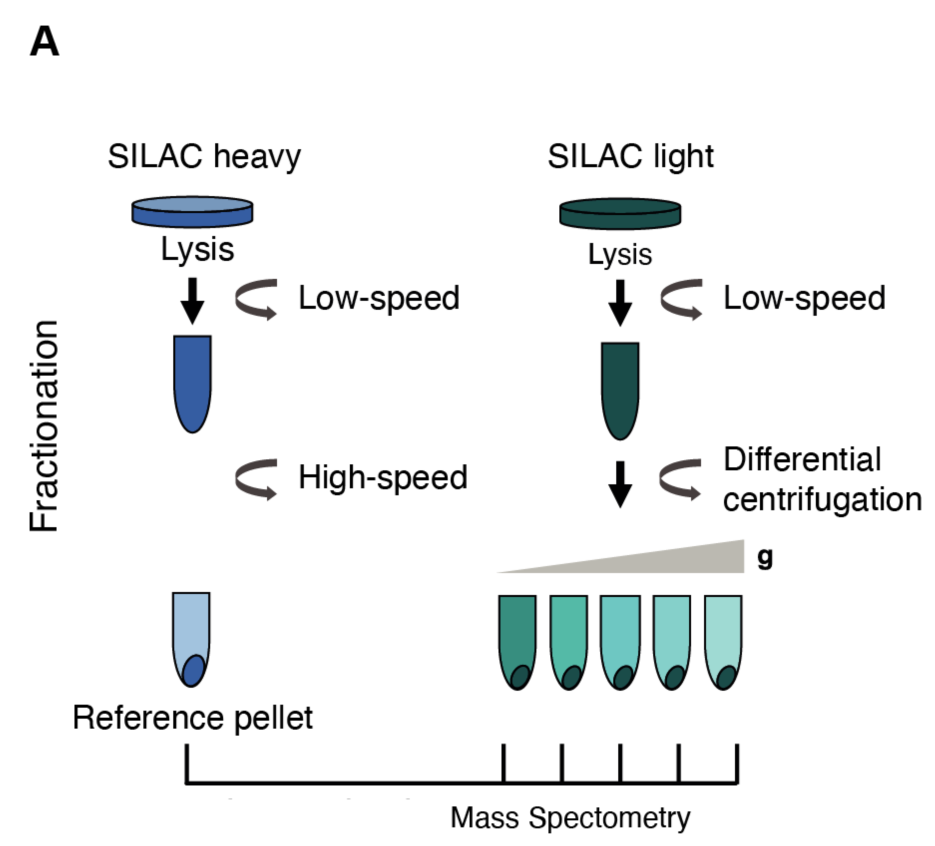
Spatial proteomics strategy to generate organellar maps. **(A)** Schematic representation of fractionation-based mass spectrometry for organellar mapping in MutuDCs. A SILAC heavy reference pellet is generated by spinning cells at a high speed and is then spiked into the light fractions. SILAC light cells are centrifuged at a range of speeds to partially separate organelles. The abundance of proteins across the fractions can then be determined by quantitative mass spectrometry.

**Fig S5.**
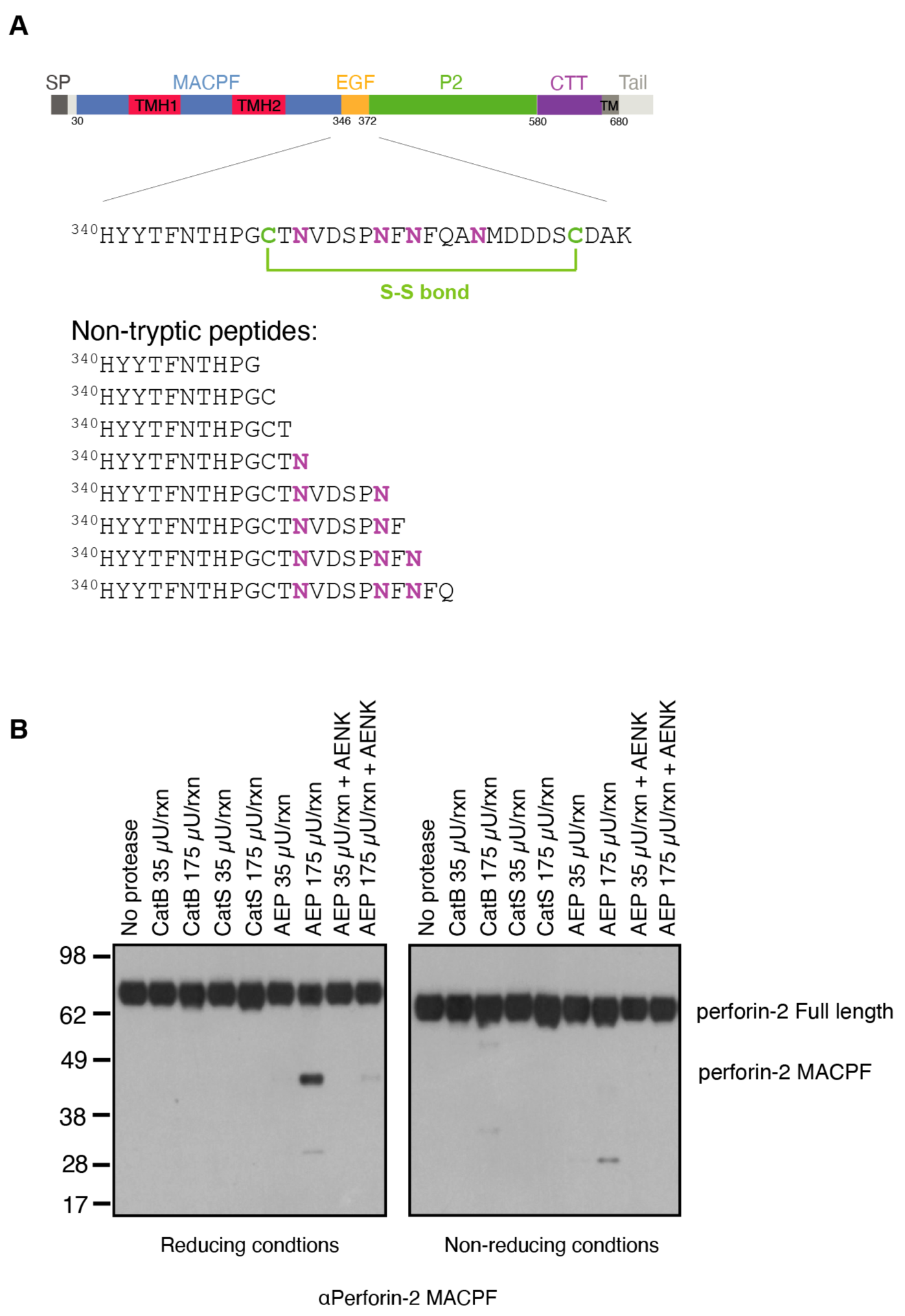
Perforin-2 in-vitro cleavage. **(A)** Schematic representation of perforin-2 highlighting the region within EGF domain encompassing the non-tryptic perforin-2 peptides detected, and the sequences of these peptides **(B)** Perforin-2 cleavage by AEP was assessed by reducing and non-reducing Western blot. AEP was preactivated prior to the cleavage reaction by incubation at 37°C for 4 h. Activated AEP was incubated with purified perforin-2 for 2 h at 37°C in the presence or absence of AEP inhibitor peptide.

**Fig S6.**
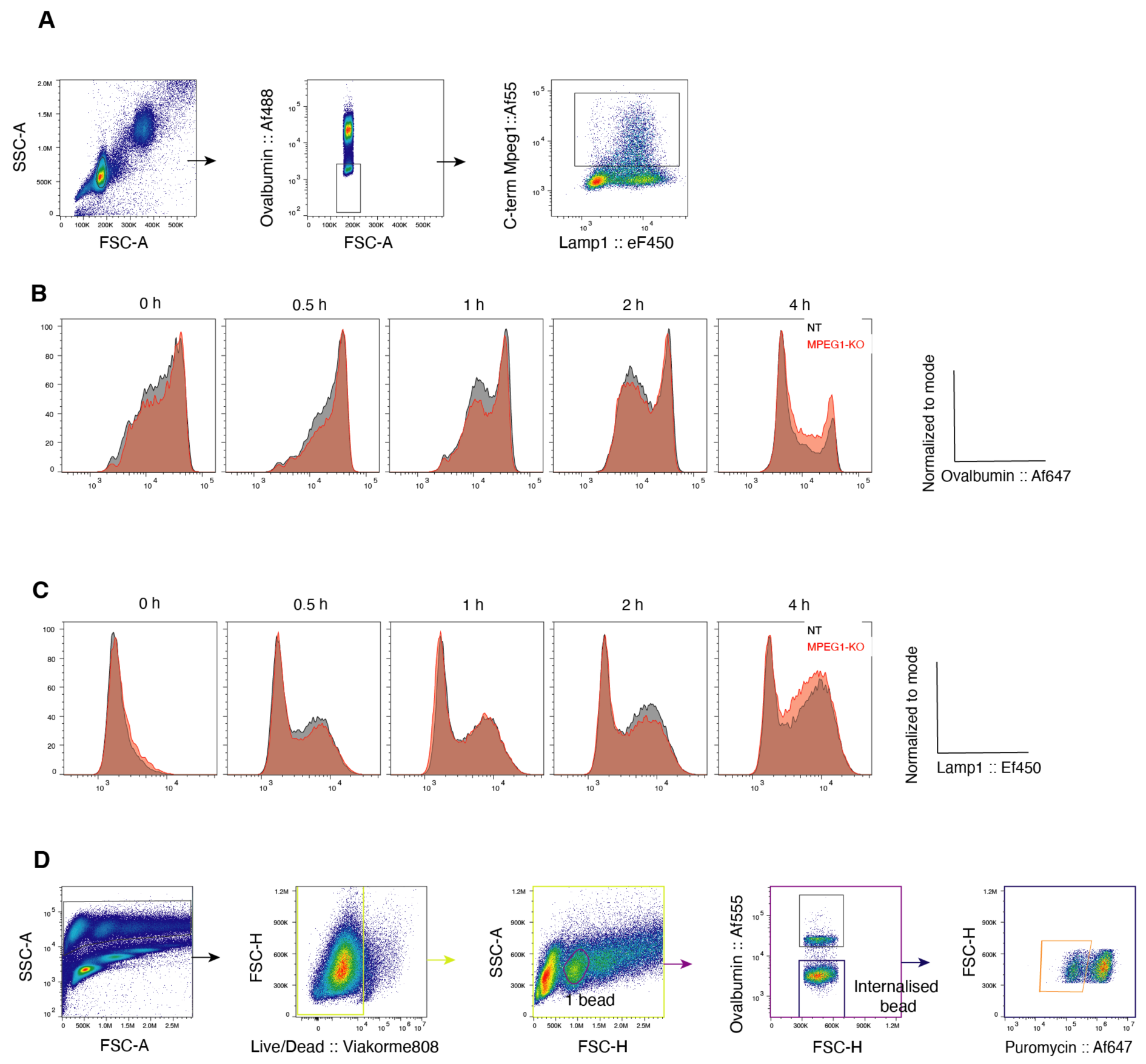
Perforin-2 does not play a role in antigen degradation or phagosome maturation. **(A)** Flow cytometry gating strategy to identify phagosomes. **(B and C)** *Mpeg1*^KO^ and NT MutuDCs were pulsed with Ova-beads and chased for the indicated times. Isolated phagosomes were stained with antibodies against (B) ovalbumin and (C) Lamp-1. Histograms are representative for three independent experiments. **(D)** Flow cytometry gating strategy to monitor translation inhibition in MutuDCs containing a single internalised bead.

**Fig S7.**
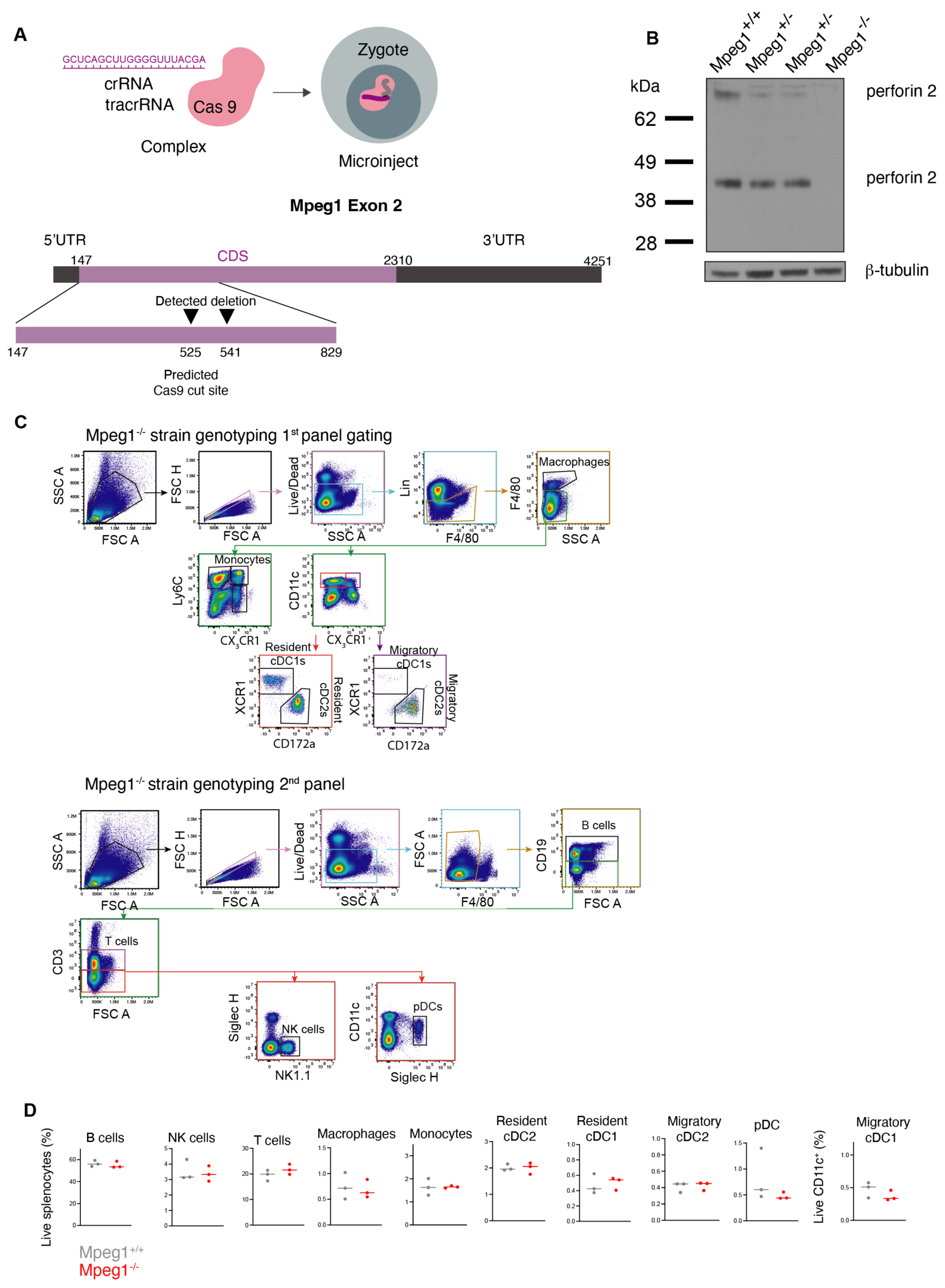
Generation and characterization of *Mpeg1-/-* mice. **(A)** CRISPR/Cas9 strategy for the generation of *Mpeg1* knock-out mice. **(B)** Perforin-2 levels in *Mpeg1*^+/+^, *Mpeg1*^+/-^ and *Mpeg1*^-/-^ splenocytes were assessed by Western blot under reducing conditions. β-tubulin was used as a loading control. **(C)** Flow cytometry gating strategy for identification of splenic immune cells in *Mpeg1*^-/-^ and *Mpeg1*^+/+^ mice. **(D)** Characterization of *Mpeg1*^-/-^ mice. Frequency of B cells, NK cells, T cells, resident and migratory cDC1s and cDC2s, macrophages, monocytes and pDCs. Data represent a single experiment using three mice per genotype.

**Fig S8.**
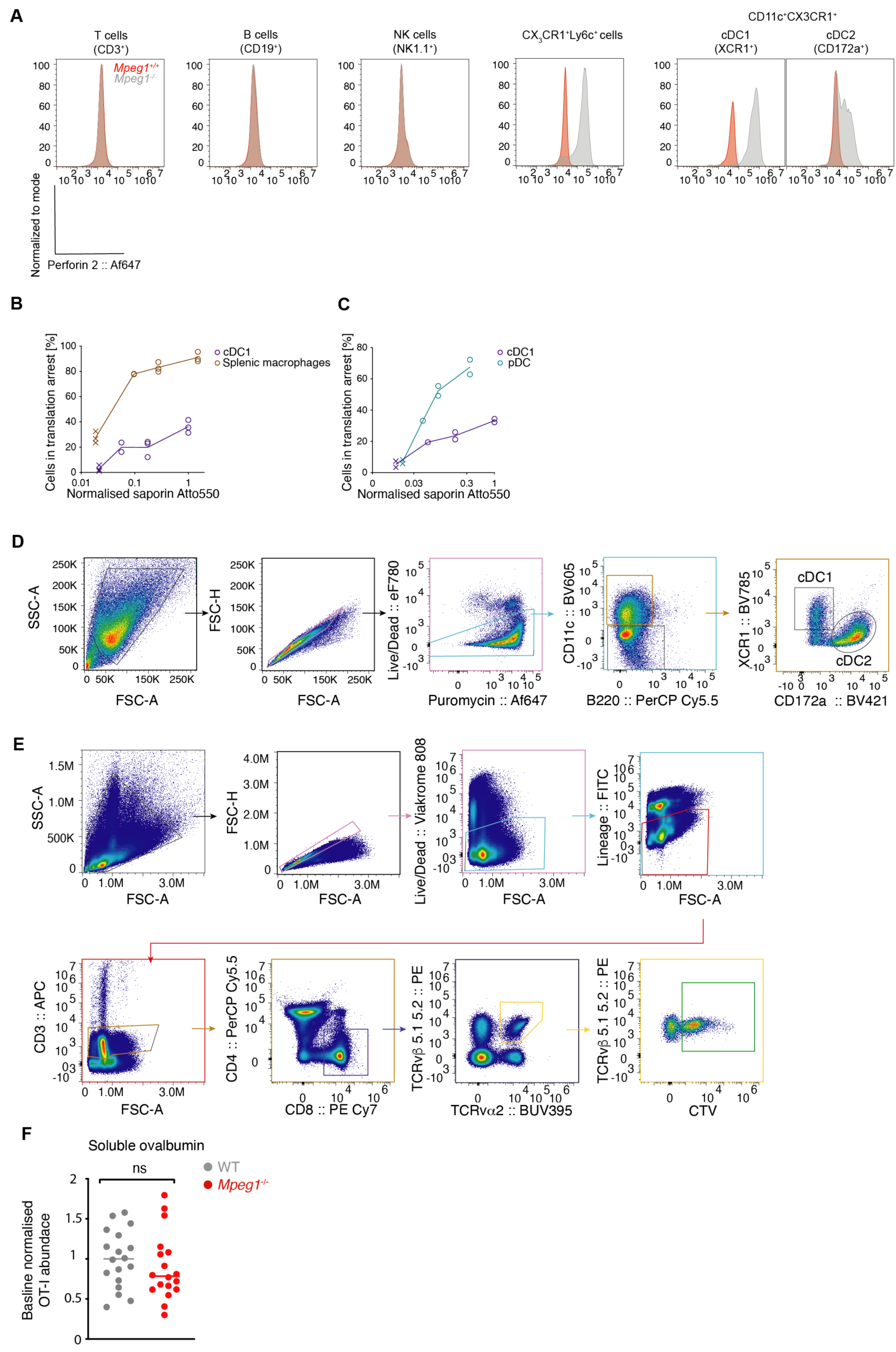
*Mpeg1-/-* mice show no defect in the cross-presentation of soluble antigens. (**A)** Perforin-2 levels in *Mpeg1*^+/+^ and *Mpeg1*^-/-^ in T cells (Live/Dead^-^, F4/80^-^, CD19^-^, CD3^-^), B cells (Live/Dead^-^, F4/80^-^, CD19^+^), NK cells (Live/Dead^-^, F4/80^-^, CD19^-^, CD3^-^, NK1.1^+^), and CD11C^+^CX3CR1^+^ cDC1s (Lineage (CD3, CD19, NK1.1^-^, F4/80^-^, CD11C^+^CX3CR1^-^, XCR1^+^) and cDC2s (Lineage (CD3, CD19, NK1.1^-^, F4/80^-^, CD11C^+^CX3CR1^-^, CD172a^+^). Histograms are representative for a single experiment using three mice per genotype. **(B and C)** CD11^+^ magnetically enriched splenocytes were pulsed with saporin (spiked 1:11 with Atto-550 labelled saporin) for 2 h, and translation was monitored by a 30 min puromycin chase. The x-axis represents Atto 550 MFI, normalised to the cDC1 Atto550 MFI at the highest saporin concentration. For the splenic macrophage/cDC1 comparison in (B), data represent three independent experiments. For the pDC/cDC1 comparison in (C), data represent two independent experiments, except for the 0.2 mg/mL concentration which was carried out once. **(D)** Flow cytometry gating strategy for Flt3L/GM-CSF cDC cultures. **(E)** Flow cytometry gating strategy for the identification of OT-I cells. **(F)** *Mpeg1*^+/+^ and *Mpeg1*^-/-^ mice were intravenously (i.v.) injected with 0.5x10^6^ CTV-labelled magnetically purified OT-I cells. One day later mice were i.v. injected with 100 μg ovalbumin and 50 μg Poly(I:C). Three days later, OT-I proliferation was assessed by flow cytometry. Data are plotted as individual values normalised to the average wild-type OT-I counts and represent five independent experiments, ns, not significant using an unpaired t-test. For gating strategy see (E).

**Table S1 Table S2 Table S3 Table S4 Table S5**

Data pertaining to the genetic screen in Fig. 2 including immgen gene expression data (cDC1s and cDC2s), a list of the sgRNAs in the custom-made library, sgRNA raw counts, CRISPR/Cas9 screen sample ID and CRISPR/Cas9 screen results.

**Table S2. Mass spectrometry of BafA1-treated MutuDCs.**

Proteomics data including protein name, normalised LFQ intensities, p-values (Student’s t-test) as well as N-term and C-term cleavage windows for BafA1-treated and control cells.

**Table S3. Table of antibodies used in this study.**

Information about the antibodies used in this study including clone, fluorochrome and dilutions.

**Table S4. Table of DNA sequences used in this study.**

Details of the plasmids, primers and gene fragments used in this study.

**Table S5. Table of reagents used in this study.**

