## Supplementary Materials for "Perforin-2 is a pore-forming effector of endocytic escape in cross-presenting dendritic cells"

### Supplementary Figures

Supp Figure 1

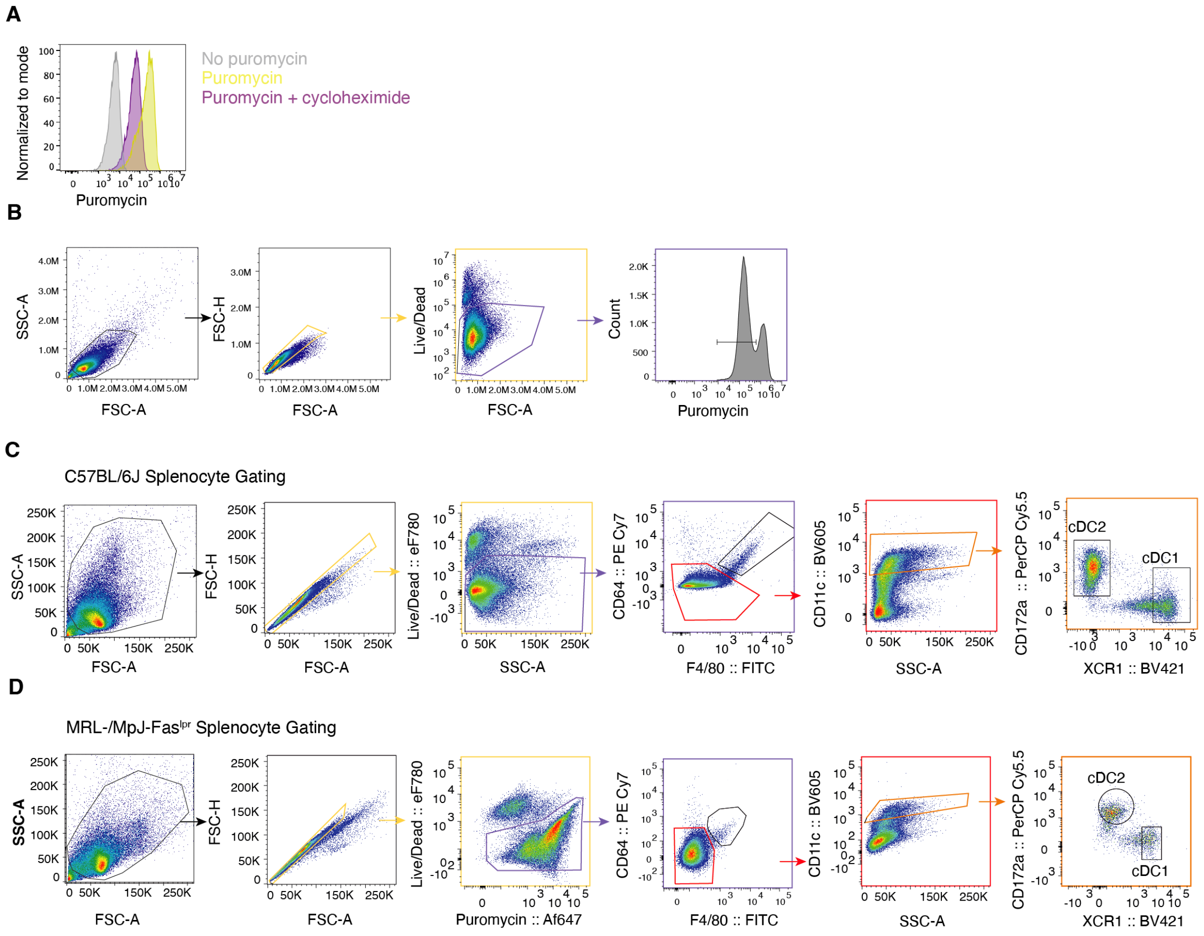


**Fig S1. Monitoring puromycin incorporation allows to track changes in translation.**
**(A)** Monitoring puromycin incorporation by flow cytometry. MutuDCs were incubated for 2 h at 37°C with 10 µg/mL cycloheximide followed by a 30 min 0.01 mg/mL puromycin chase in the presence or absence of 10 µg/mL cycloheximide. Puromycin incorporation was monitored with a αPuromycin antibody. Histograms are representative for three independent experiments.

**(B)** Flow cytometry gating strategy to monitor translation inhibition in MutuDCs.

**(C)** Flow cytometry gating strategy for C57BL/6J splenic cDC1s and cDC2s for the *ex vivo* saporin assay.

**(D)** Flow cytometry gating strategy for MRL/MpJ-Fas$^{lpr}$ splenic cDC1s and cDC2s for the *ex vivo* saporin assay.

Supp Figure 2

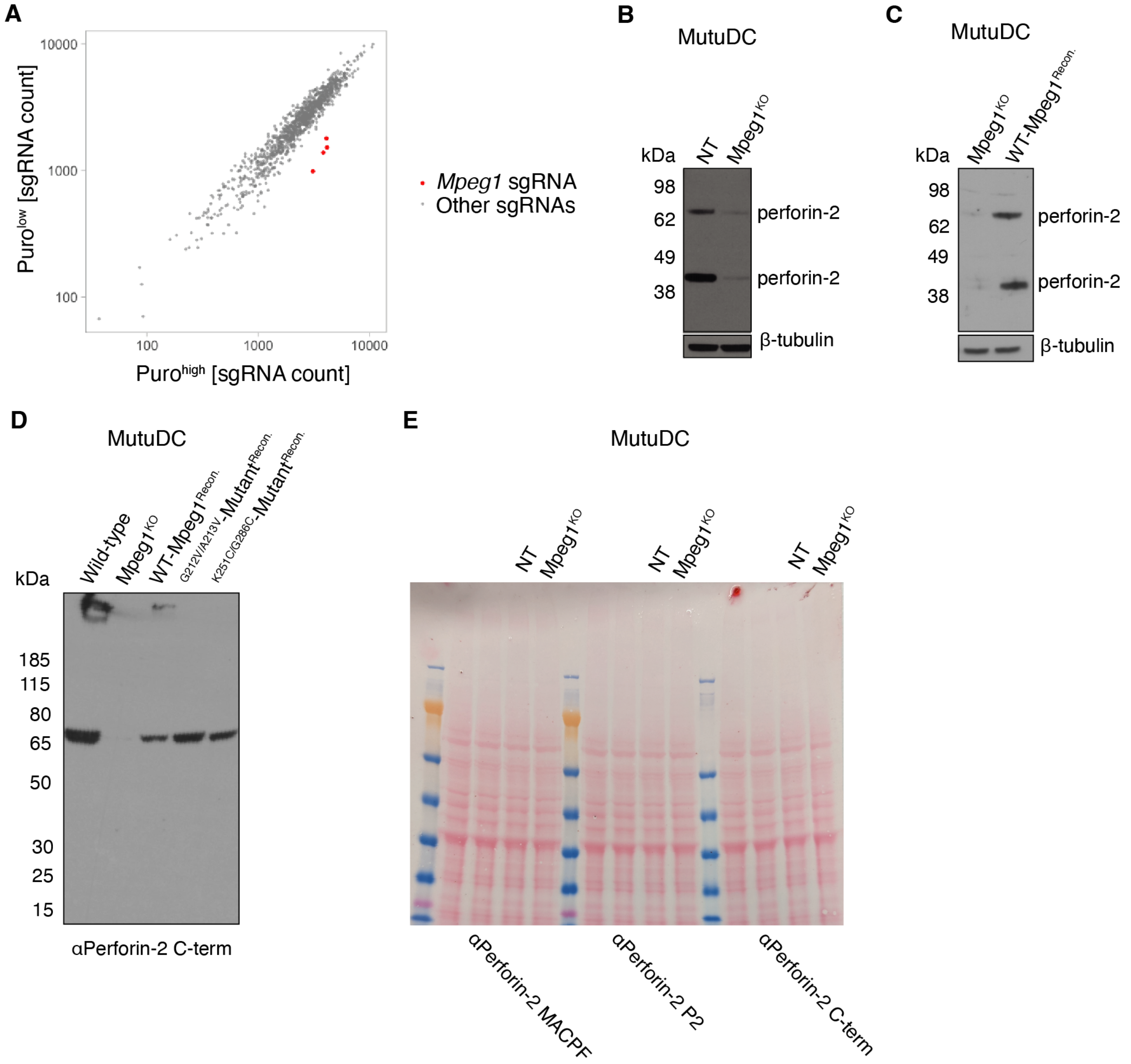


**Fig S2. Identification of *Mpeg1* in the CRISPR-Cas9 genetic screen and characterization of the *Mpeg1* KO and rescue MutuDC lines.**

**(A)** Relative abundance of sgRNAs in puro$^{high}$ and puro$^{low}$ populations from the saporin-puromycin-based genetic screen. The screen was performed in three biological repeats. The sgRNA counts from each population in each screen replicate were normalised, and the average of the counts for puro$^{high}$ and puro$^{low}$ populations is plotted. Each dot corresponds to one sgRNA. The sgRNAs targeting *Mpeg1* are highlighted in red.

**(A)** Perforin-2 protein levels in *Mpeg1*$^{KO}$ and NT MutuDCs were assessed by Western blot under reducing conditions. β-tubulin was used as a loading control.

**(B)** Perforin-2 protein levels in *Mpeg1*$^{KO}$ and *Mpeg1*$^{KO}$ MutuDCs complemented with sgRNA resistant *Mpeg1* were assessed by Western blot under reducing conditions. $\beta$-tubulin was used as a loading control.

**(C)** Protein levels of WT and perforin-2 mutants in reconstituted *Mpeg1*$^{KO}$ MutuDCs were assessed by Western blot under non-reducing conditions.

**(D)** Ponceau staining corresponding to the Western blot in Fig. 5C.

Supp Figure 3

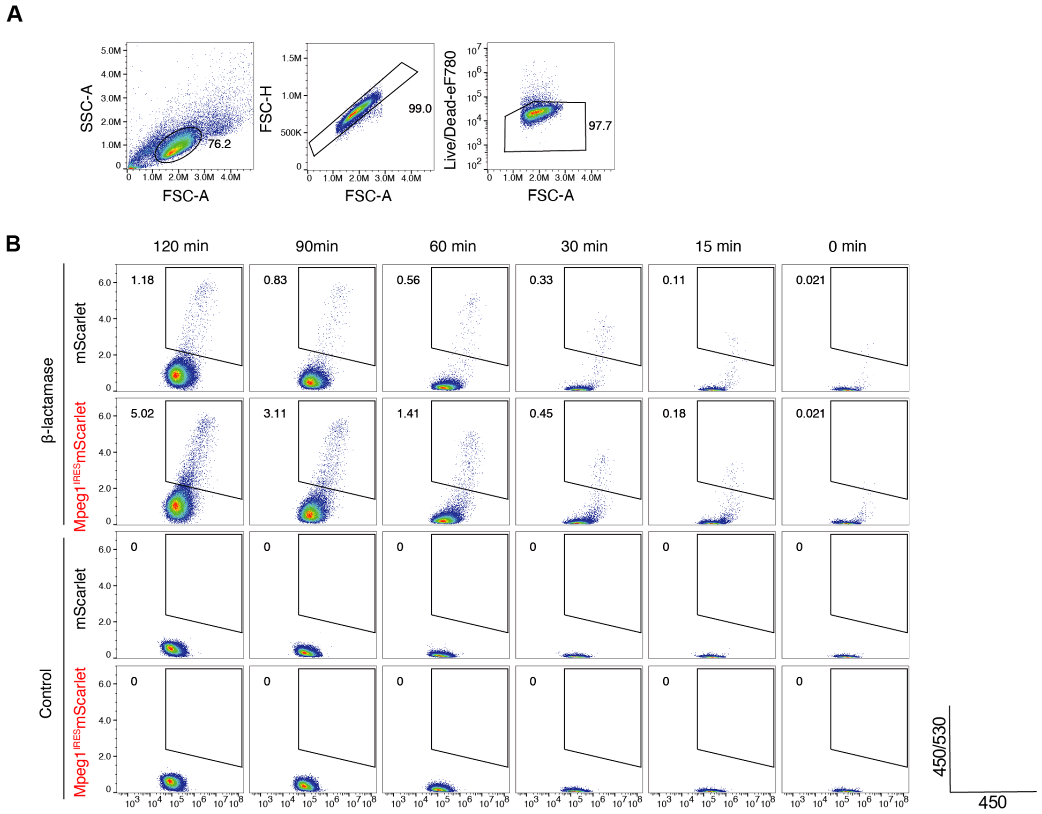


**Fig S3. Gating strategy for the CCF4 β-lactamase assay.**

**(A)** Flow cytometry gating strategy for the CCF4 β-lactamase assay in mScarlet and Mpeg1[IRES]-mScarlet HeLa cells.

**(B)** Representative gating for CCF4 cleavage for each condition. Gates were defined to exclude spontaneous conversion of CCF4 in the absence of β-lactamase over time.

Supp Figure 4

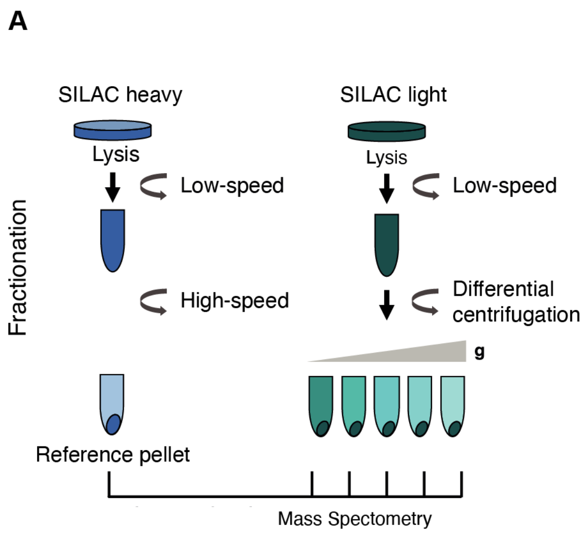


**Fig S4. Spatial proteomics strategy to generate organellar maps.**
**(A)** Schematic representation of fractionation-based mass spectrometry for organellar mapping in MutuDCs. A SILAC heavy reference pellet is generated by spinning cells at a high speed and is then spiked into the light fractions. SILAC light cells are centrifuged at a range of speeds to partially separate organelles. The abundance of proteins across the fractions can then be determined by quantitative mass spectrometry.

Supp Figure 5

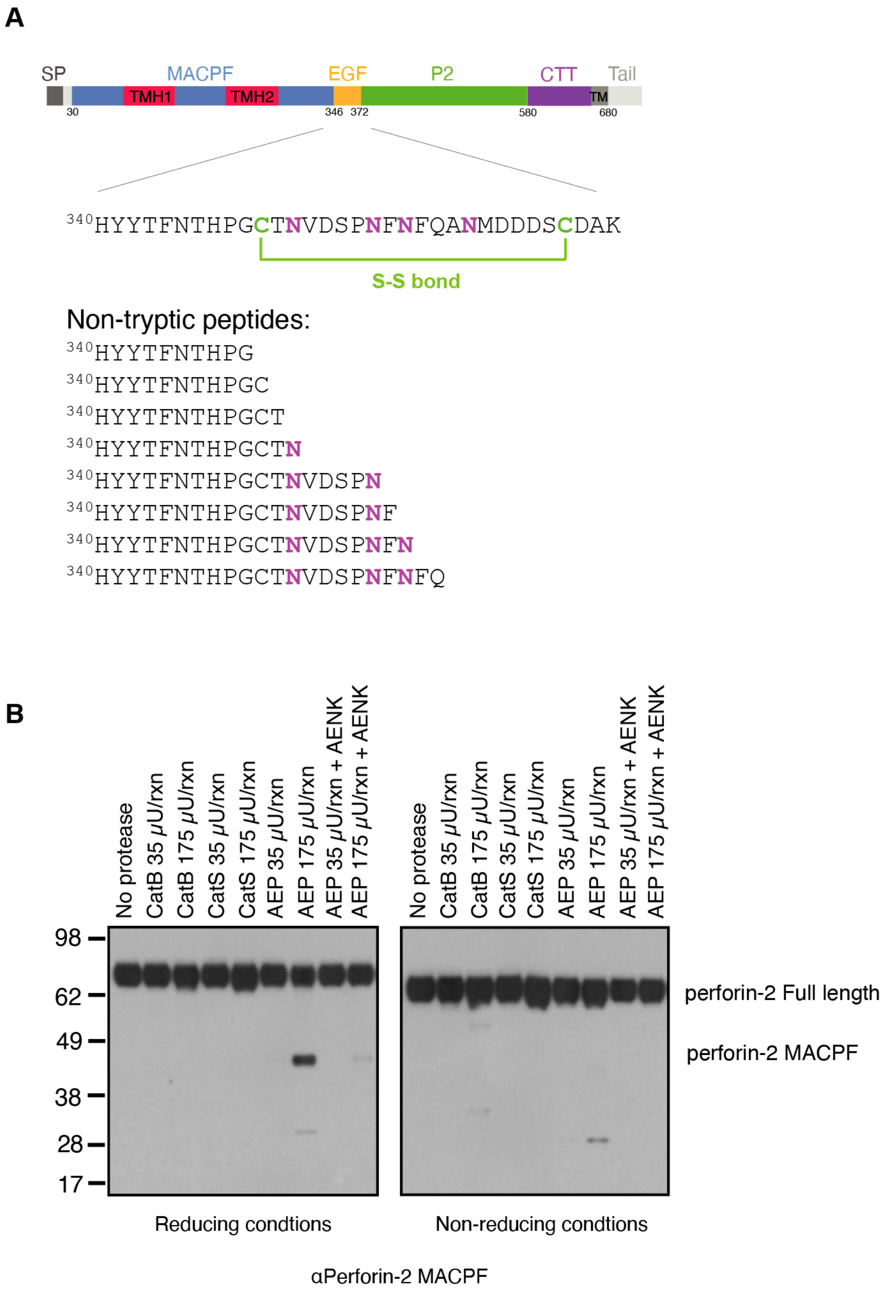


**Fig S5. Perforin-2 in-vitro cleavage.**

**(A)** Schematic representation of perforin-2 highlighting the region within EGF domain encompassing the non-tryptic perforin-2 peptides detected, and the sequences of these peptides

**(B)** Perforin-2 cleavage by AEP was assessed by reducing and non-reducing Western blot. AEP was preactivated prior to the cleavage reaction by incubation at 37°C for 4 h. Activated AEP was incubated with purified perforin-2 for 2 h at 37°C in the presence or absence of AEP inhibitor peptide.

Supp Figure 6

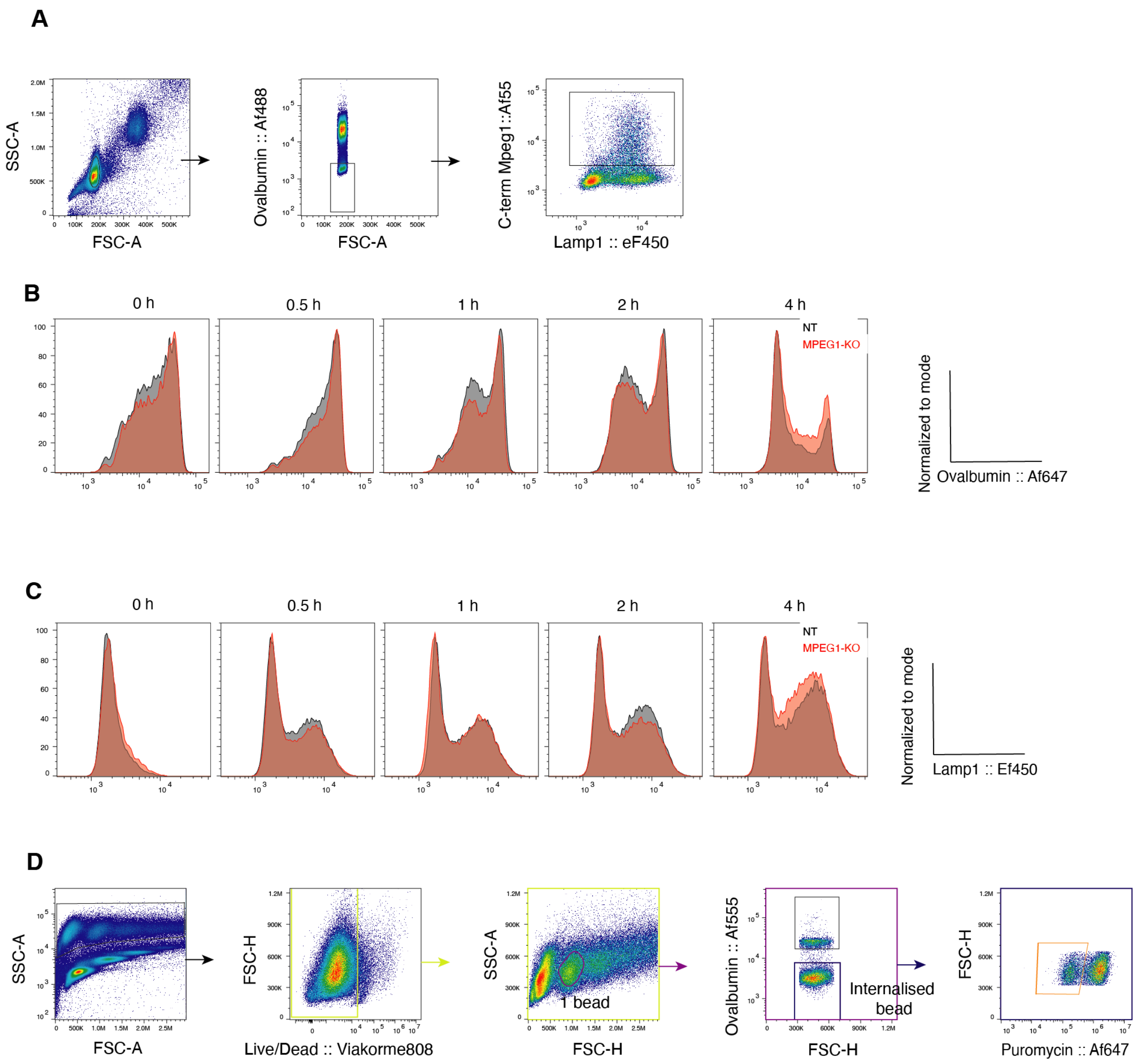


**Fig S6. Perforin-2 does not play a role in antigen degradation or phagosome maturation.** **(A)** Flow cytometry gating strategy to identify phagosomes.

**(B and C)** *Mpeg1*$^{KO}$ and NT MutuDCs were pulsed with Ova-beads and chased for the indicated times. Isolated phagosomes were stained with antibodies against (B) ovalbumin and (C) Lamp-1. Histograms are representative for three independent experiments.

**(D)** Flow cytometry gating strategy to monitor translation inhibition in MutuDCs containing a single internalised bead.

Supp Figure 7

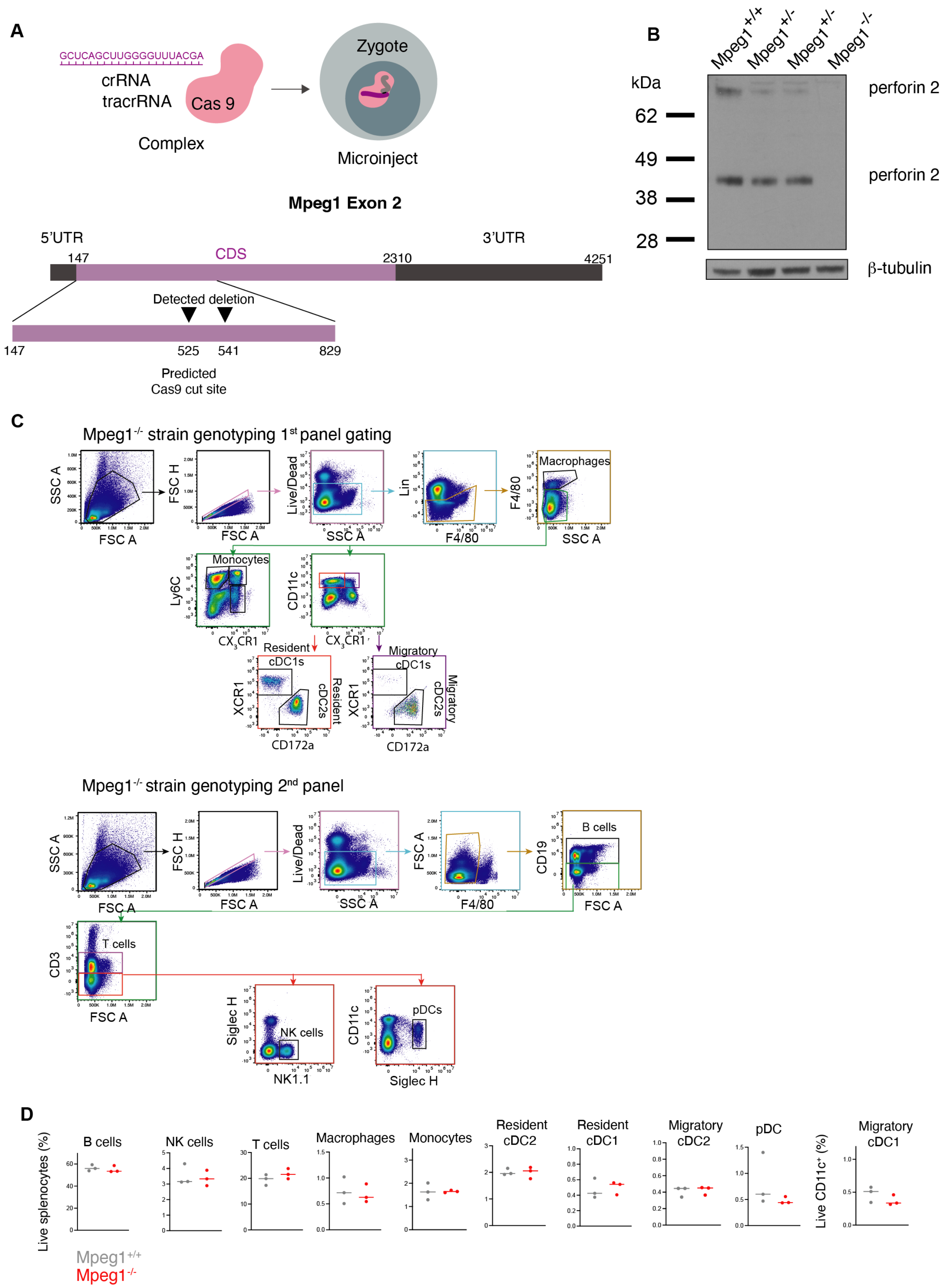


480

**Fig S7. Generation and characterization of *Mpeg1*$^{-/-}$ mice.**

**(A)** CRISPR/Cas9 strategy for the generation of *Mpeg1* knock-out mice.

**(B)** Perforin-2 levels in *Mpeg1*$^{+/+}$, *Mpeg1*$^{+/-}$ and *Mpeg1*$^{-/-}$ splenocytes were assessed by Western blot under reducing conditions. β-tubulin was used as a loading control.

**(C)** Flow cytometry gating strategy for identification of splenic immune cells in *Mpeg1*$^{-/-}$ and *Mpeg1*$^{+/+}$ mice.

**(D)** Characterization of *Mpeg1*$^{-/-}$ mice. Frequency of B cells, NK cells, T cells, resident and migratory cDC1s and cDC2s, macrophages, monocytes and pDCs. Data represent a single experiment using three mice per genotype.

Supp Figure 8

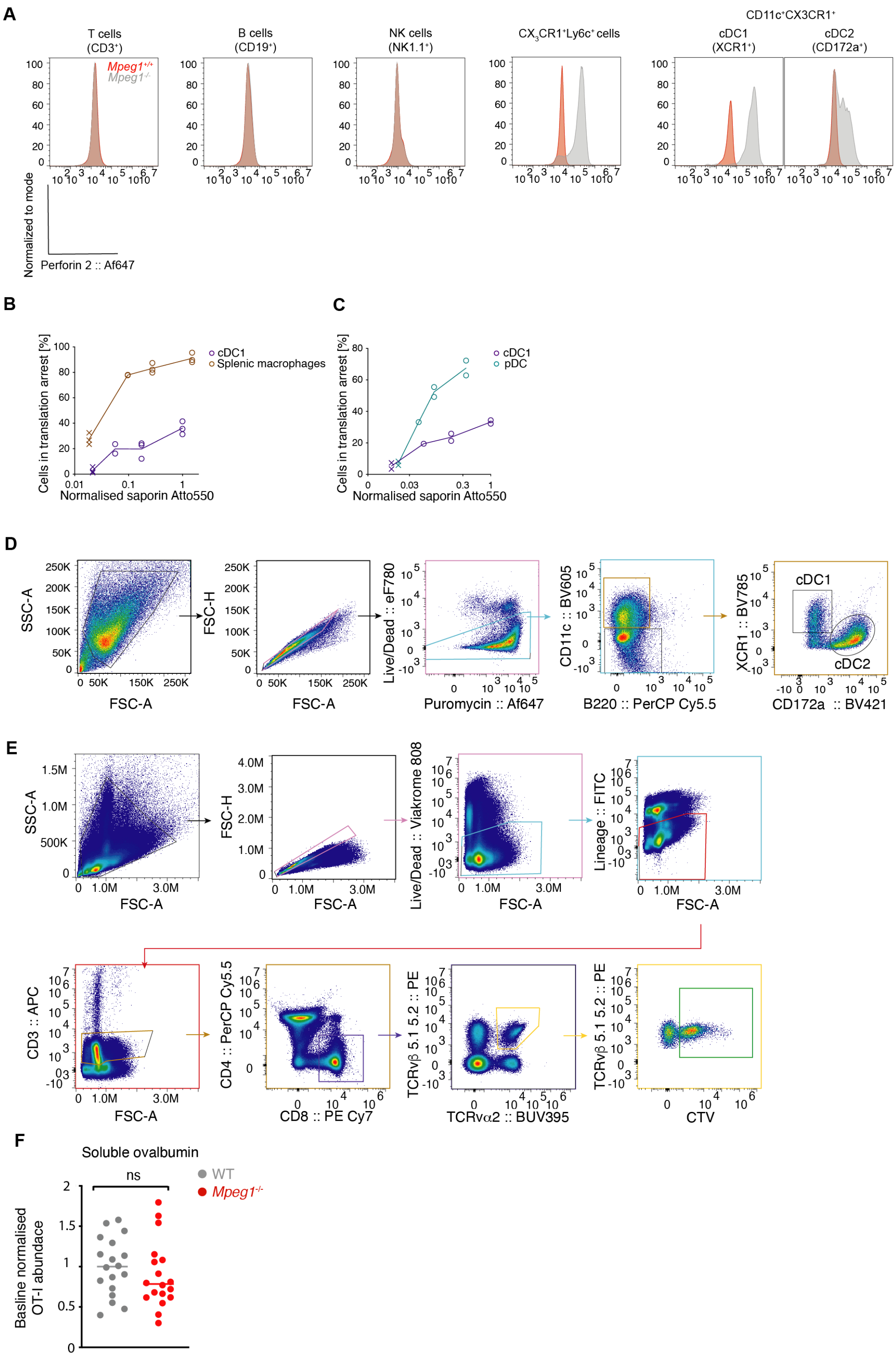


**Fig S8. *Mpeg1$^{-/-}$* mice show no defect in the cross-presentation of soluble antigens.**
(**A**) Perforin-2 levels in *Mpeg1$^{+/+}$* and *Mpeg1$^{-/-}$* in T cells (Live/Dead$^-$, F4/80$^-$, CD19$^-$, CD3$^-$), B cells (Live/Dead$^-$, F4/80$^-$, CD19$^+$), NK cells (Live/Dead$^-$, F4/80$^-$, CD19$^-$, CD3$^-$, NK1.1$^+$), and CD11C$^+$CX$_3$CR1$^+$ cDC1s (Lineage (CD3, CD19, NK1.1$^-$, F4/80$^-$, CD11C$^+$CX$_3$CR1$^-$, XCR1$^+$) and

cDC2s (Lineage (CD3, CD19, NK1.1$^-$, F4/80$^-$, CD11C$^+$CX$_3$CR1$^-$, CD172a$^+$). Histograms are representative for a single experiment using three mice per genotype.
